# Knowledge Gaps of STIs in Africa; Systematic review

**DOI:** 10.1101/557389

**Authors:** M M Badawi, M A Salah-Eldin, A B Idris, E A Hasabo, Z H Osman, W M Osman

## Abstract

Sexually Transmitted Infections (STIs) are ambiguous burden of tremendous health, social and economic consequences, The current systematic review was conducted in order to determine awareness and knowledge of Africans of sexually transmitted infections, not only concerning HIV/AIDS, but also other STIs such as, gonorrhea, syphilis, HBV, HCV and HPV. A systematic review of the literature was conducted, studies were retrieved and selected after they fulfilled the inclusion criteria and passed the assessment procedure. related data was extracted, quantitative analysis was conducted among participants who responded to questions related to HIV, HBV, HCV, HPV or STIs knowledge, sensitivity analysis as well as subgroup analysis were also conducted. Seventy four articles addressing knowledge among 35 African countries were included and 136 questions were analyzed and synthesized. The question Using condom will reduce HIV transmission?” was answered by 1,799,374 Africans in 35 countries, 66.82% [95% Cl; 62.65, 70.98] answered yes. While the question “Is sexual contact a possible route of HBV transmission?” was answered by 7,490 participants in 5 countries; 42.58% [95% Cl; 20.45, 64.71] answered yes. The differences observed among populations are highlighting the possibility for containment and control by directing light toward specific populations or countries as well as addressing specific awareness knowledge to ensure that the general as well as the related specific preventive awareness knowledge is improved.

## Introduction

Sexually transmitted Infections (STIs) are ambiguous burden of tremendous health, social and economic consequences. Most of them are hidden because many people may feel stigmatized when addressing them. Moreover, the committee on prevention and control of sexually transmitted diseases in USA estimated that the annual costs of selected major STDs are approximately $10 billion or, if sexually transmitted HIV is included, $17 billion ^[1]^.

According to UNAIDS; almost 37 million people globally were living with HIV in 2017, sub-Saharan Africa accounted for 66% of the cases, 68% of new adult HIV infections, 92% of new infections in children and 72% of all AIDS-related deaths. Earlier in 2009, Swaziland topped the world’s HIV epidemic countries with a 26% prevalence among adults, while South Africa was the country with the world’s largest prevalence of people living with HIV as 5.6 million ^[2,3]^.

On the other hand and according to WHO; an estimated 257 million people are living with HBV infection with the highest prevalence in the Western Pacific Region and the African Region as 6.2% and 6.1% of the adult population are infected, respectively. About 1% of persons living with HBV infection (2.7 million people) are also infected with HIV. Moreover, approximately 399,000 people die each year from hepatitis C infection and the estimated global HPV prevalence is 11.7% with the Sub-Saharan Africa having the largest burden as well (24.0%), ^[4–6]^.

Africa is considered the continent with the lowest Gross Domestic Product (GDP) as most African countries fall within the lower-middle to low income countries classification. Moreover, In March 2013, despite of the predicted uprising in African economy in the following decades, Africa was identified as the world’s poorest inhabited continent; Africa’s entire combined GDP is estimated to be barely a third of the United States’, this could be straightforwardly influencing screening opportunities, medical consultations as well as treatment options. Taking that under consideration; a strategy of STIs containment in Africa should primarily emphasize prevention and its related knowledge. Chan and Tsai in their study represented awareness outcome of data collected from 33 sub-Saharan African countries. Although their study determined the estimated awareness according to data collected from 2003 to 2015 as well as a knowledge trend among each participated country was illustrated, awareness of five questions were assessed regarding HIV only. The current systematic review was conducted in order to determine awareness and knowledge of Africans of sexually transmitted infections, not only concerning HIV/AIDS, but also other STIs such as, gonorrhea, syphilis, HBV, HCV and HPV and concerning all awareness determinants that are reported in the literature ^[7,8]^.

## Materials and methods

### Search strategy

To identify relevant studies, a systematic review of the literature was conducted in the 1^st^ of December 2018. The review was conducted in accordance with the PRISMA (Preferred Reporting Items for Systematic Reviews and Meta-Analyses) Statement ^[8]^ (Table S1). A comprehensive search was conducted in PubMed, Embase, Google scholar, Scopus, Index Copernicus, DOAJ, EBSCO-CINAHL, Cochrane databases without language limits (studies written in French were later excluded). To obtain a current situation evidence; only studies published in or after 2010 were included. Furthermore, all studies where the data collection process took place before 2010 were also excluded, the only exception was if the collection process continued months/years earlier than 2010 and ended in 2010 or afterwards. The keywords used in PubMed was as follow:

((HIV[Tiab] OR syphilis[Tiab] OR gonorrhea[Tiab] OR sexual behavior[Tiab] OR “men who have sex with men”[Tiab] OR condom[Tiab] OR “herpes simplex virus”[Tiab] OR “sex workers”[Tiab] OR sex [tiab]OR human immunodeficiency virus[Tiab] OR HBV[Tiab] OR HCV[Tiab] OR HPV[Tiab] OR prostitutes[Tiab] OR trichomonas vaginalis[Tiab]) AND (behavior [Ti] OR risk [ti] OR awareness[Ti] OR knowledge[Ti] OR assessment[Ti]) AND (africa[Tiab] OR algeria[Tiab] OR angola[Tiab] OR benin[Tiab] OR botswana[Tiab] OR burkinafaso[Tiab] OR burundi[Tiab] OR caboverde[Tiab] OR cameroon[Tiab] OR central african republic[Tiab] OR CAR[Tiab] OR chad[Tiab] OR comoros[Tiab] OR “democratic republic of the congo”[Tiab] OR “republic of the congo”[Tiab] OR cote d’ivoire[Tiab] OR djibouti[Tiab] OR egypt[Tiab] OR equatorial guinea[Tiab] OR eritrea[Tiab] OR eswatini[Tiab] OR swaziland[Tiab] OR ethiopia[Tiab] OR gabon[Tiab] OR gambia[Tiab] OR ghana[Tiab] OR guinea[Tiab] OR guinea-bissau[Tiab] OR kenya[Tiab] OR lesotho[Tiab] OR liberia[Tiab] OR libya[Tiab] OR madagascar[Tiab] OR malawi[Tiab] OR mali[Tiab] OR mauritania[Tiab] OR mauritius[Tiab] OR morocco[Tiab] OR mozambique[Tiab] OR namibia[Tiab] OR niger[Tiab] OR nigeria[Tiab] OR rwanda[Tiab] OR (sao tome principe[Tiab] OR senegal[Tiab] OR seychelles[Tiab] OR sierra leone[Tiab] OR somalia[Tiab] OR south africa[Tiab] OR south sudan[Tiab] OR sudan[Tiab] OR swaziland[Tiab] OR eswatini[Tiab] OR tanzania[Tiab] OR togo[Tiab] OR tunisia[Tiab] OR uganda[Tiab] OR zambia[Tiab] OR zimbabwe[Tiab])).

Moreover, to optimize our search, hand searches of reference lists of included articles were also performed.

### Study selection and data extraction

All authors independently assessed titles and abstracts for eligibility, and any disagreement was resolved through discussion. A copy of the full text was obtained for all research articles that were available and approved in principle to be included. Abstraction of data was in accordance with the task separation method; method and result sections in each study were separately abstracted in different occasions to reduce bias ^[9]^. Moreover, data abstraction was conducted with no consideration of author’s qualifications or expertise. Studies assessed the knowledge of parasitic infections as well as studies conducted among healthcare workers (clinicians, laboratory specialists, nurses, dentists and midwives) were excluded. If a data regarding the period of conduction is missing in a study the reference list was crossed, if any cited study is published after 2010 authors of the current review agreed to predict that the study is conducted after 2010 and hence it was considered for inclusion and it was considered to be addressed later in the review as (conducted after 2010), otherwise the study was excluded. All studies measuring awareness level with scores or if it is generally good or moderate or bad without determining further details were excluded. Each research article was screened for all relevant information and recorded in the data extraction file (Microsoft Excel), as one article may report outcome of awareness and/or knowledge and/or attitude toward specific sexually transmitted infection or toward several STIs, in a single population or among several ones. Moreover, data from each method section was extracted using a predefined set of variables; study characteristics, type of participants, study population size, geographical region and the period of the study conduction.

### Assessment of quality

Each included article was evaluated based on a framework for making a summary assessment of the quality. The related published literature was crossed, then a framework was structured specifically to determine the level of representativeness of the studied population and to judge the strength of the estimates provided. Six questions were to be answered in each article, each answer represent either 1 score for yes, 0 score for No or 0 score for not available; A total score for risk of bias and quality was calculated by adding up the scores in all six domains, resulting in a score of between 0 and 6. The highest score indicates the highest quality, studies with a score for quality greater or equal to 3 (higher quality) were included in the review.

The six domains were: is the study objective clearly defined?, is the study sample completely determined?, is the study population clearly defined and specified?, is the response rate of participants above 70%?, is the methodology used rigorous? and is the data analysis rigorous?

Trim and Fill method was used to assess the risk of publication bias in each question responses in the included studies ^[10]^. Publication bias was assessed separately for each question-corresponding responses only if the question was addressed and answered in studies equal or greater than ten.

### Quantitative analysis

Meta-analysis was performed - whenever possible using Review Manager Software (Version 5.3). In studies where the Standard Error (SE) is not reported, the following formula was used to calculate it: SE√p=(1-p)/ n where p stands for Prevalence. The software automatically provided the Confidence Interval (CI) according to the calculated SE, if the CI is provided in a study; it was introduced accordingly. The heterogeneity of each meta-analysis was assessed, the random effects was favored over the fixed effects model in all meta-analysis established as differences between studies is predicted to be possible due to the diversity of the study populations. Sensitivity analysis was also approached to determine the effect of studies conducted in populations proposed to behave in indifference manners or proposed to be more aware on the overall pooled prevalence. Moreover, subgroup analysis was also conducted -whenever suitable to determine awareness level in specific country or population. A question to take part in the meta-analysis has to be included in at least two studies. Moreover, for providing the full picture as well as emphasizing potential research gaps; all HIV-related questions that are proposed to be of interest according to the objective of the current review and was answered by at least 1,000 Africans but included only in one study were also provided alongside their related references, however, questions related to other STIs were provided regardless of the number of participants due to their minority. Questions with similar outcome were proposed to be the same (i.e. the question “do you think sexual intercourse will increase the risk of HIV transmission?” and the question “is HIV sexually transmitted?” were considered as one question.

## Results

### Studies included

A total of 7,540 articles were identified from the search strategy including hand searches of reference lists of included original research articles and reviews. From these, 7,453 articles were excluded. Seventy four articles met our inclusion criteria and passed the quality assessment procedure. The articles reported specific awareness determinants and/or knowledge and/or attitudes of an African population regarding STIs as general and/or HBV and/or HCV and/or HPV and/or HIV. (Fig. 1) illustrates the PRISMA flow diagram. The included articles are depicted in (Table 1). Assessment of the quality of included studies is depicted in (Table S2).

**Table (1):**
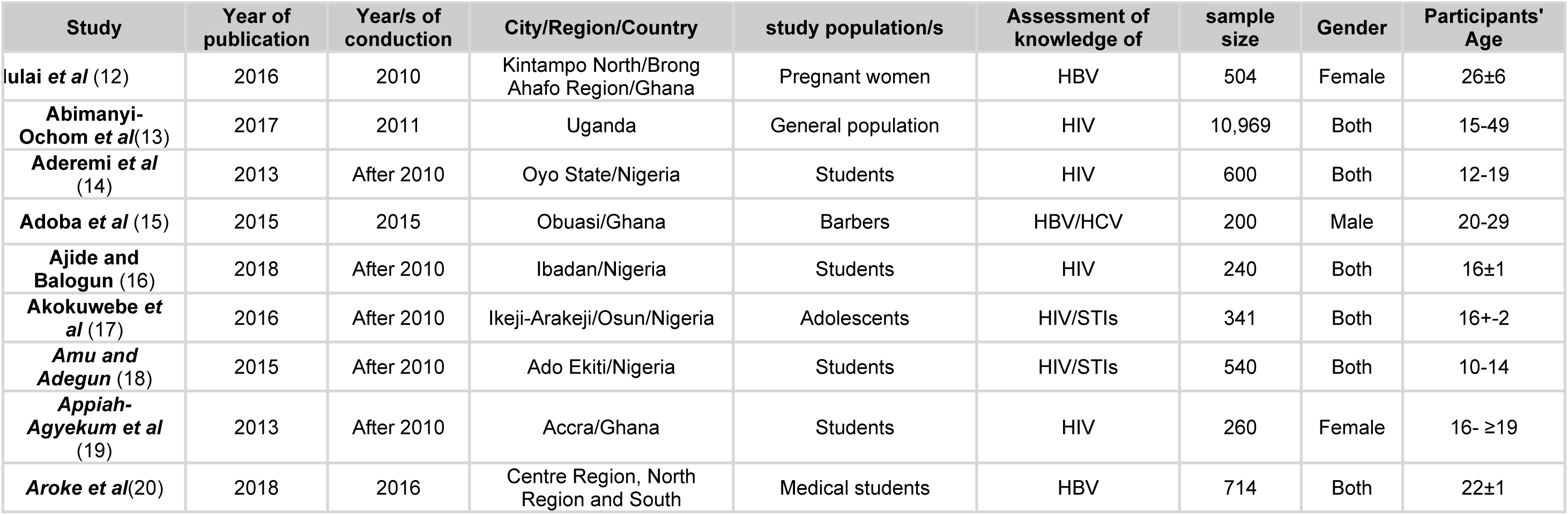
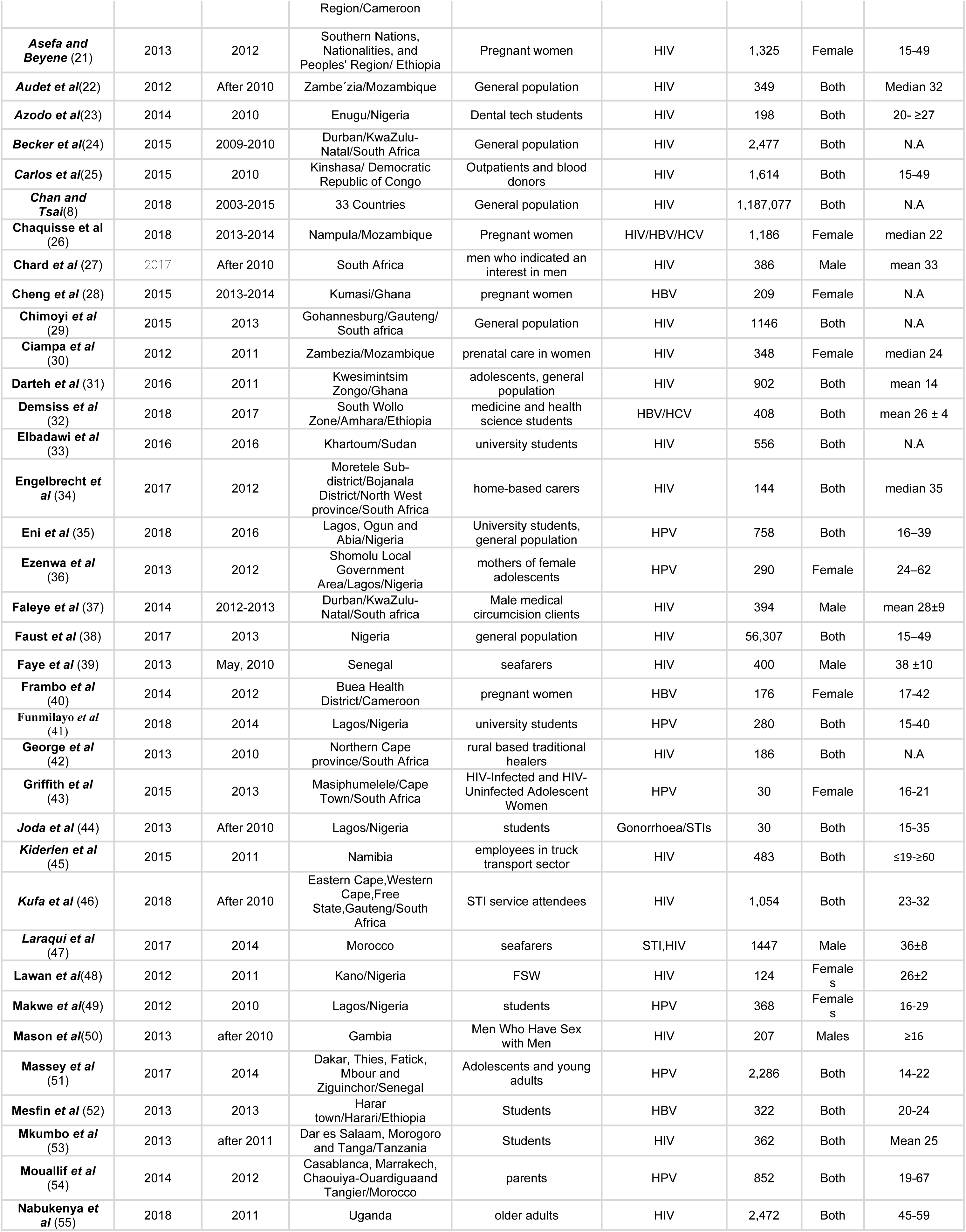
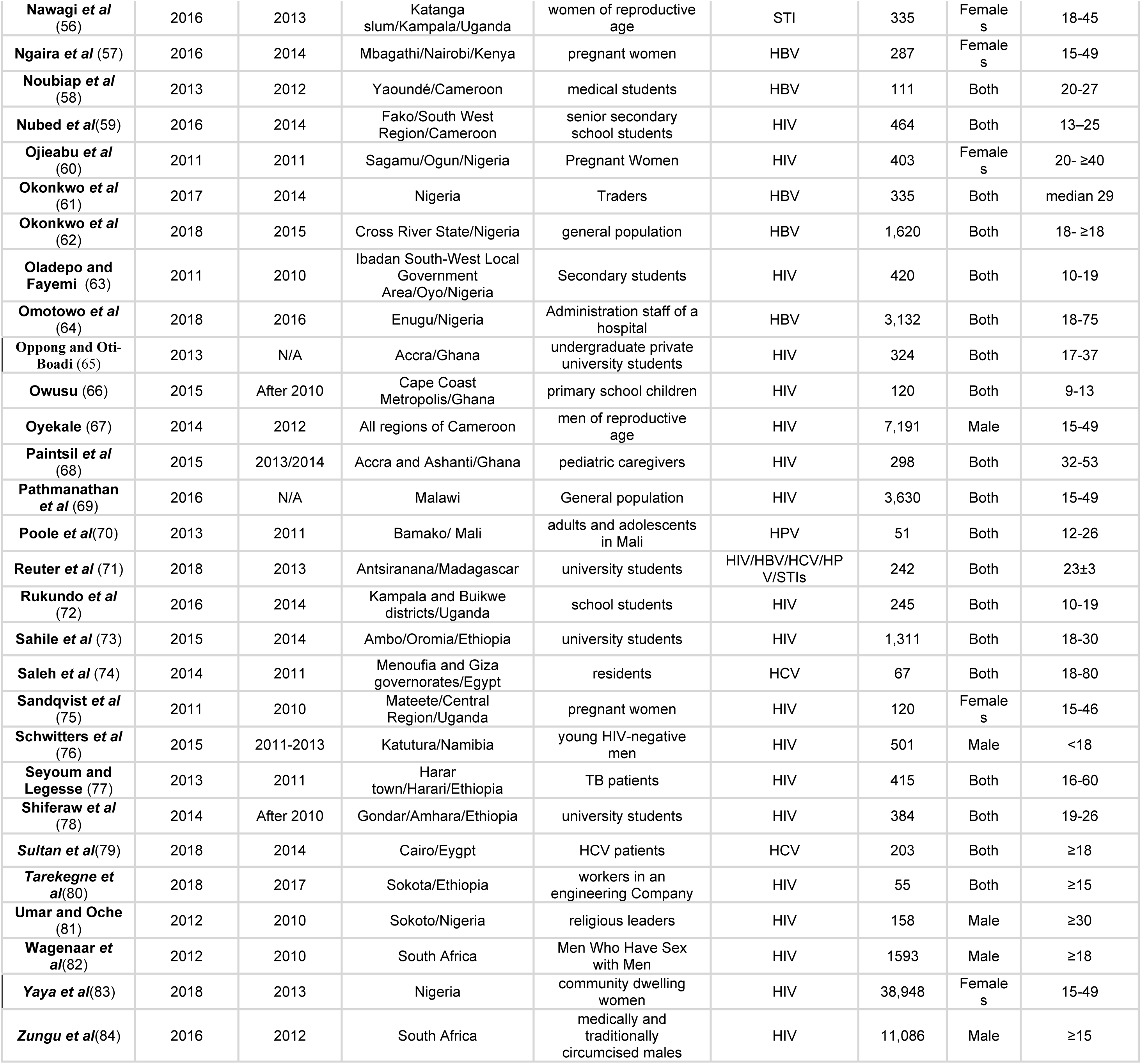
Characteristics of included studies

**Fig (1):**
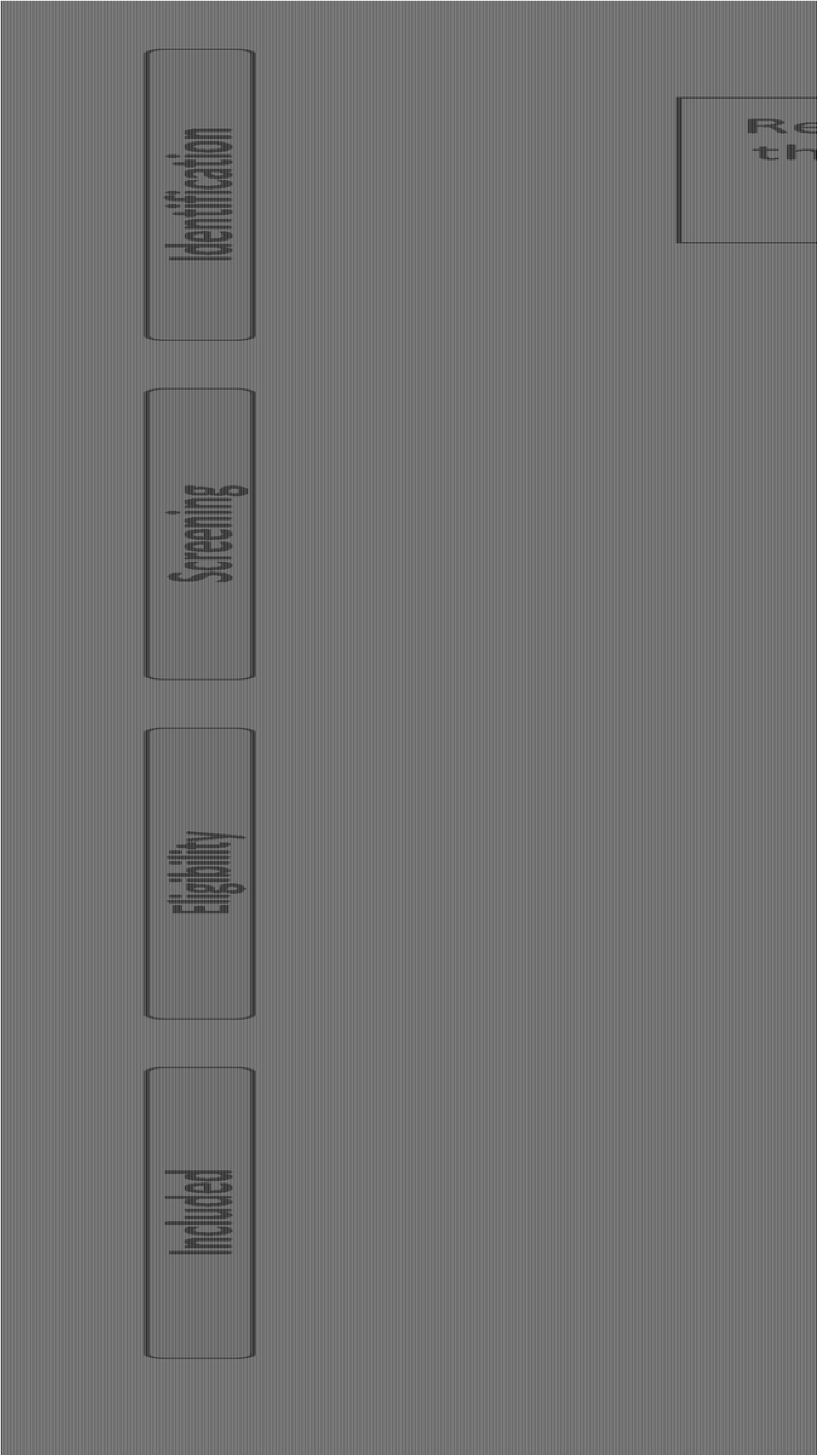
Literature search and selection of studies (PRISMA flow diagram).

### Study characteristics

The characteristics of the included studies are depicted in (Table1), among which the oldest were published in 2010 while the most recent ones were published in 2018. Fifty one research articles determining HIV awareness level and/or knowledge and/or attitudes were included, while 14 articles determining HBV awareness level and/or knowledge and/or attitudes were included. Furthermore, 6 and 9 articles concerned of awareness level and/or knowledge and/or attitudes level regarding HCV and HPV were included, respectively. Seven articles determining STIs awareness level and/or knowledge and/or attitudes as general were also included. Two hundred questions were summarized among which 136 questions were analyzed and synthesized from included studies including the subgroup analysis. Publication bias assessment indicated no major asymmetry (data not shown).

### Human Immunodeficiency Virus (HIV)

Fifty one included studies assessed the awareness of 1,235,811 Africans in regard to HIV in total of 35 countries, eleven studies were conducted in Nigeria ^[14,16,83,17,18,23,38,48,60,63,81]^, nine in South Africa ^[24,27,29,34,37,42,46,82,84]^, five in each of Ghana ^[19,31,65,66,68]^ and Ethiopia ^[21,73,77,78,80]^, four in Uganda ^[13,55,72,75]^, three in Mozambique ^[22,26,30]^, two in each of Namibia ^[45,76]^ and Cameroon ^[59,67]^, one in each of Congo ^[25]^, Sudan ^[33]^, Senegal ^[39]^, Morocco ^[47]^, Gambia ^[50]^, Tanzania ^[53]^, Madagascar ^[71]^ and Egypt ^[74]^ while a study provided awareness prevalence in 33 countries ^[8]^. The conduction of the studies ranged from 2010 to 2017. Population under study was distributed among students and adolescents, general population, pregnant women, female sex workers, male sex workers or males who show interest of males, TB patients, seafarers and other occupations (Table 1). Majority of studies were conducted among both genders (34/51), eight studies were toward females only while nine were toward males only. Age of respondents ranged from 10 to 60 years (Table 1). Forty two questions were asked to the participants that are related to the knowledge and awareness of HIV as general, transmission routes, clinical symptoms, pathological consequences and prevention attitude, among which 31 questions were analyzed and synthesized. The question “ Using condom will reduce HIV transmission?” was answered by 1,799,374 Africans in Benin, Burkina Faso, Burundi, Cameroon, Chad, Comoros, Cote d’Ivoire, Democratic Republic of Congo, Ethiopia, Gabon, Ghana, Guinea, Kenya, Lesotho, Liberia, Madagascar, Malawi, Mali, Morocco, Mozambique, Namibia, Niger, Nigeria, Rwanda, São Tomé and Príncipe, Senegal, Sierra Leone, South Africa, Sudan, Swaziland, Tanzania, Togo, Uganda, Zambia and Zimbabwe; 66.82% [95% Cl; 62.65, 70.98] answered yes. The question ‘’ Is HIV contracted through Sexual intercourse?’’ was answered by 252,482 participants in South Africa, Ethiopia, Uganda, Madagascar, Ghana, Nigeria, Gambia, Morocco, Namibia, Senegal, Sudan and Mozambique; 72.20% [95% Cl; 64.25, 80.16] answered yes. Questions asked, their corresponding articles’ data, the pooled prevalence, the pooled prevalence after conducting sensitivity analysis and the confidence intervals are depicted in (Table 2 & Fig 2). Heterogeneity was high in all questions (I^2^ more than 80%), except for the question “ Is TB associated with HIV infection?” where I^2^ = 0%.

**Table (2):**
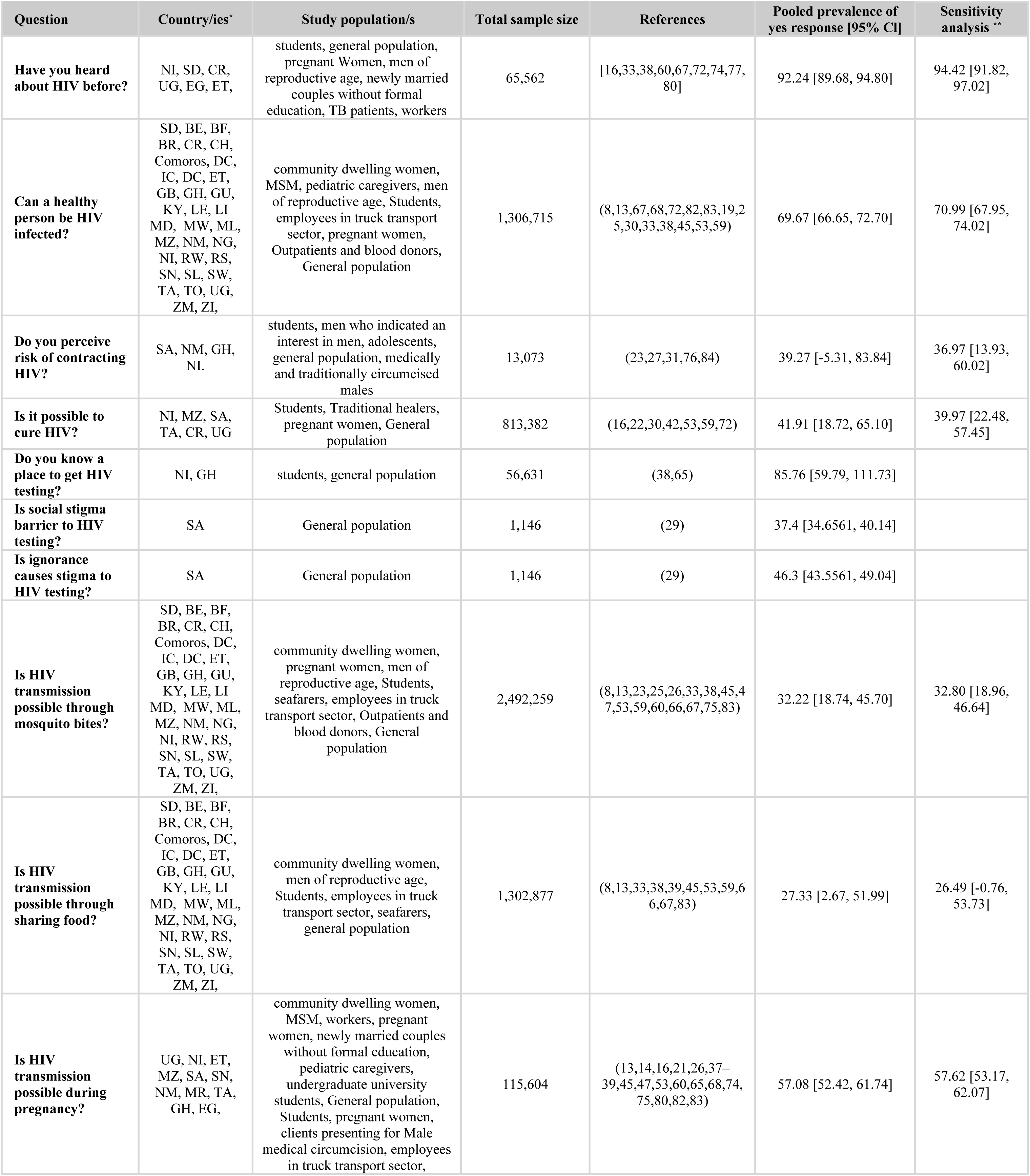
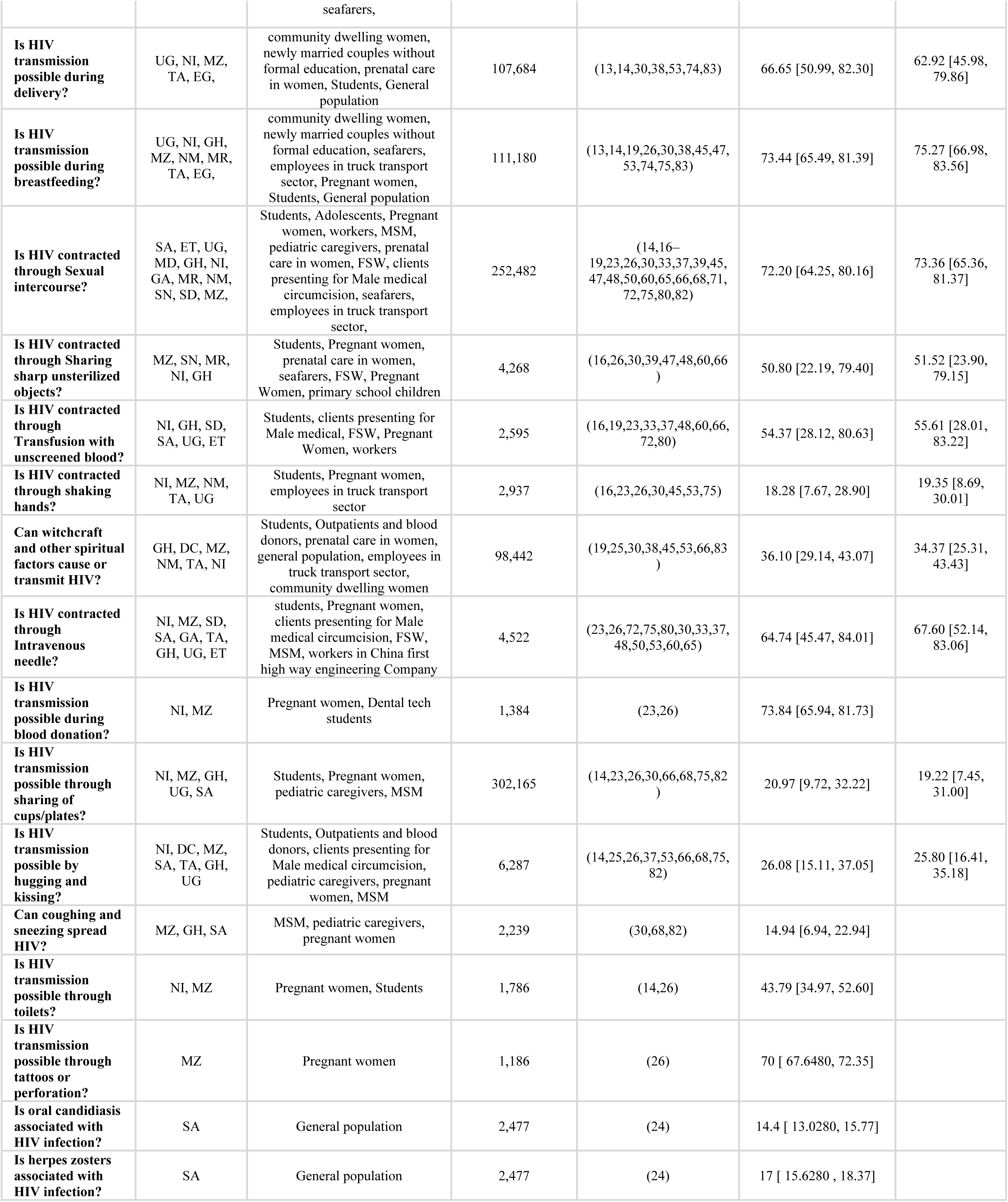
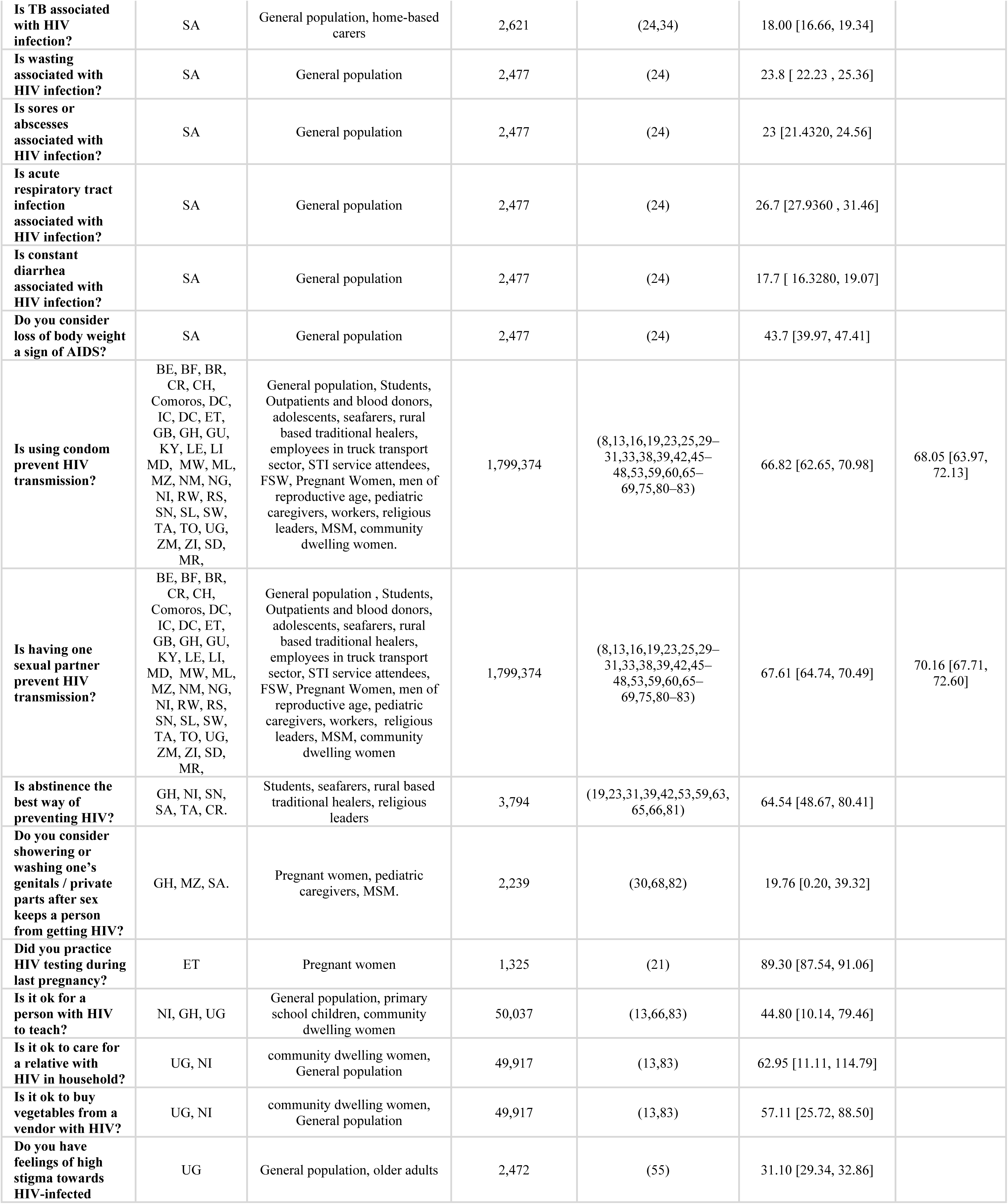
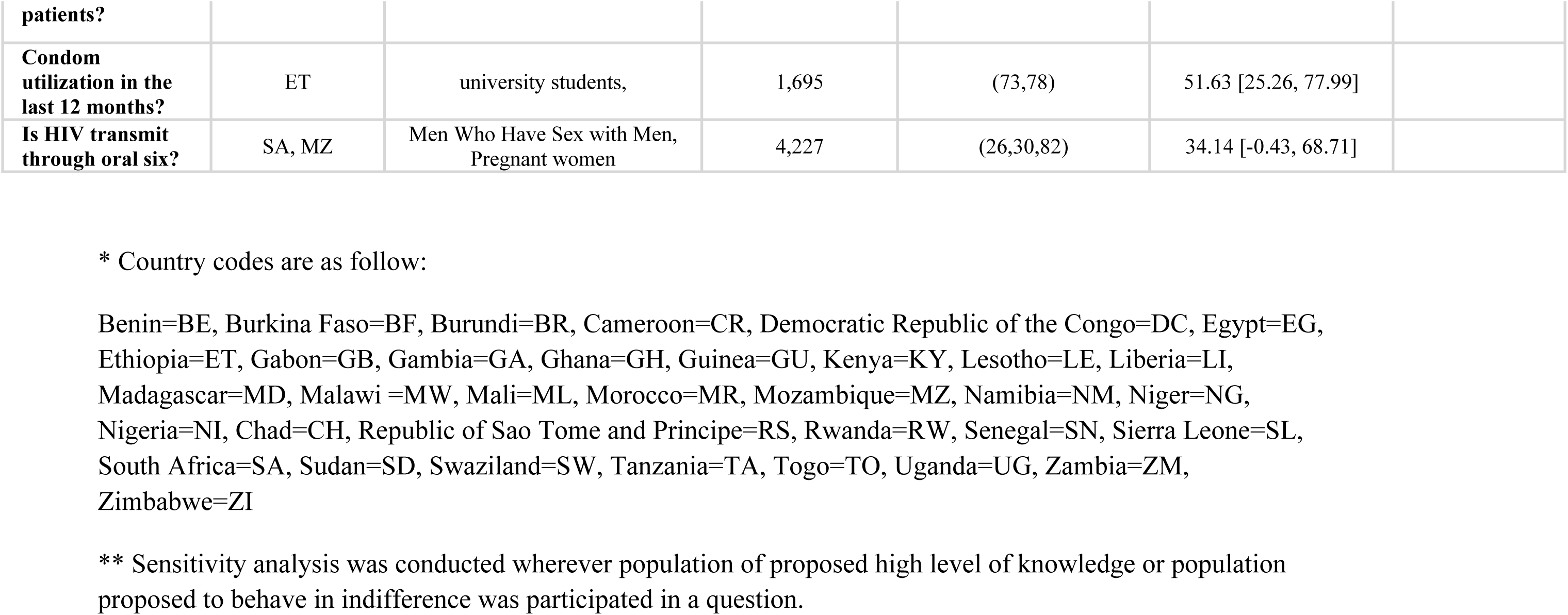
Awareness of HIV-related knowledge among Africans

**Fig (2):**
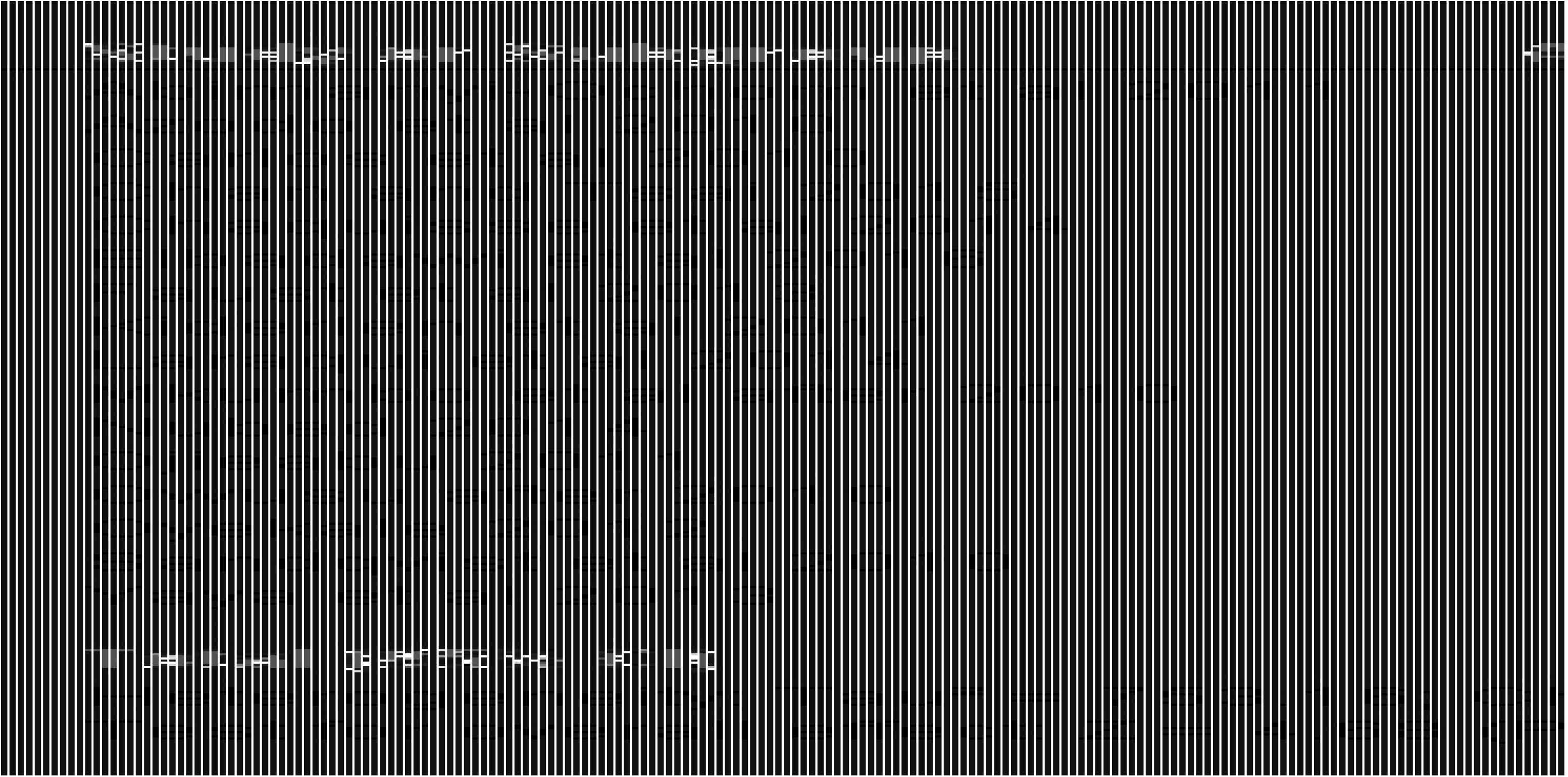
Meta analysis of 1,799,374 Africans’ yes response to the question “Is using condom prevent HIV transmission?”

### Nigeria

Eleven included studies in regard to HIV were conducted among Nigerians representing a total population of 98,279 participants; three studies were conducted in Oyo State ^[14,16,63]^ and one in each of Osun State ^[17]^, Ekiti State ^[18]^, Enugu State ^[23]^, Kano State ^[48]^, Sokoto State ^[81]^ and Ogun State ^[60]^. Two studies were nationally representative and participants were from different States ^[38,83]^. The oldest among the study included were conducted in 2010 while the newest were conducted in 2013 (Table S3). Population under study was found to be mainly students and adolescents (6/11), while one was toward each of pregnant women, religious leaders, general population, female sex workers and community dwelling women (Table S3). Majority of studies were conducted among both genders (7/11), three studies were toward females only while one study included only males. Age of respondents ranged from 12 to 49 years. Twenty two questions were asked to the participants that are related to the knowledge and awareness of HIV as general, transmission routes, clinical symptoms, pathological consequences and prevention attitude, among which 18 questions were analyzed and synthesized. The question ‘’ Using condom will reduce HIV transmission?’’ was answered by 57,430 participants; 52,67% [95% Cl; 44.42, 60.91] answered yes. The question ‘Is HIV can be transmitted through mosquito bites?’’ was answered by 95,856 participants; 16.86% [95% Cl; 6.77, 26.95] answered yes. Questions asked, their corresponding study’s characteristics, the pooled prevalence and the confidence intervals are depicted in (Table 3). Heterogeneity was high in all questions (I^2^ more than 80%).

**Table (3):**
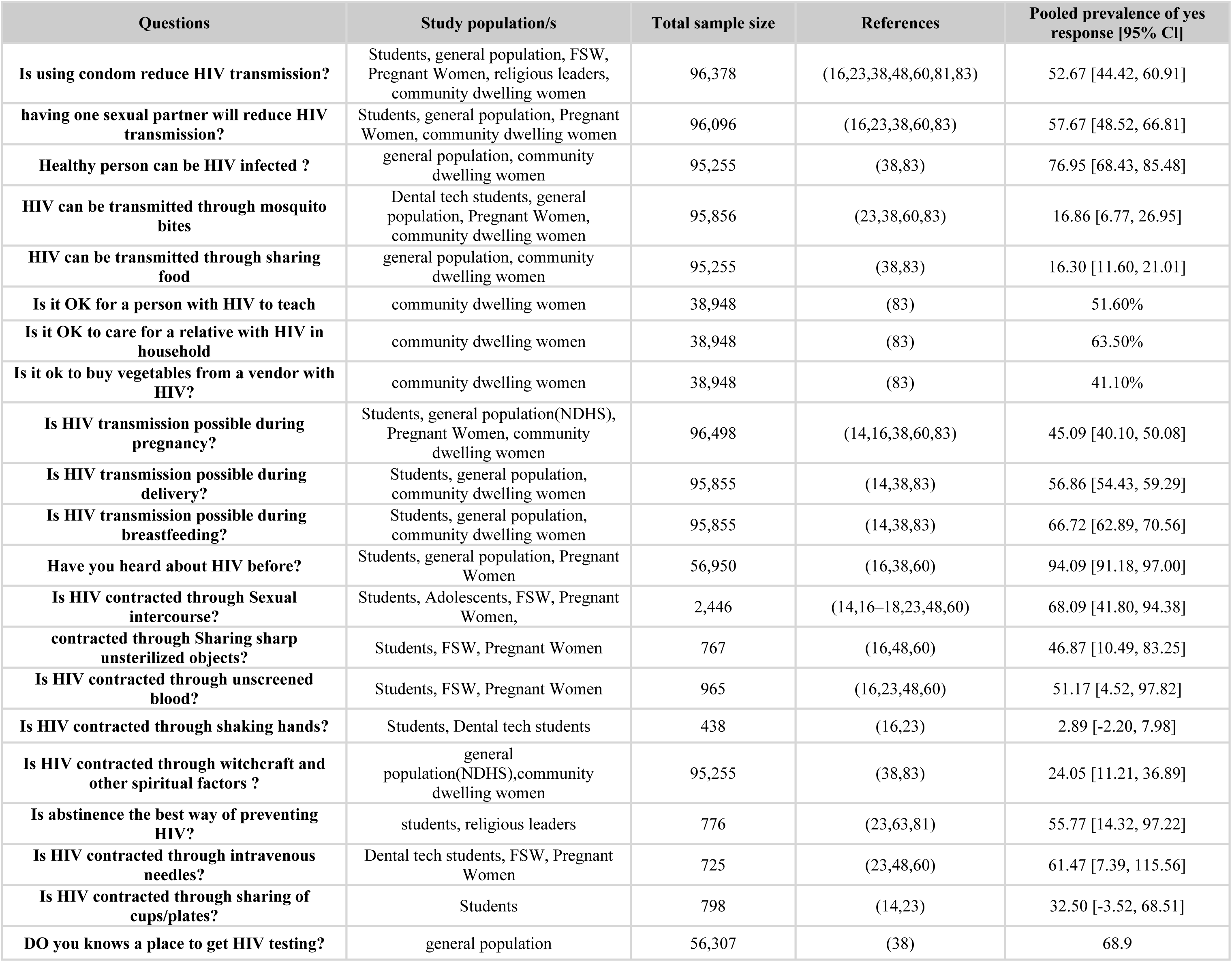
Awareness of HIV-related knowledge among Nigerian population

| Questions | Study population/s | Total sample size | References | Pooled prevalence of yes response [95% CI] |
| --- | --- | --- | --- | --- |
| Is using condom reduce HIV transmission? | Students, general population, FSW, Pregnant Women, religious leaders, community dwelling women | 96,378 | (16,23,38,48,60,81,83) | 52.67 [44.42, 60.91] |
| having one sexual partner will reduce HIV transmission? | Students, general population, Pregnant Women, community dwelling women | 96,096 | (16,23,38,60,83) | 57.67 [48.52, 66.81] |
| Healthy person can be HIV infected ? | general population, community dwelling women | 95,255 | (38,83) | 76.95 [68.43, 85.48] |
| HIV can be transmitted through mosquito bites | Dental tech students, general population, Pregnant Women, community dwelling women | 95,856 | (23,38,60,83) | 16.86 [6.77, 26.95] |
| HIV can be transmitted through sharing food | general population, community dwelling women | 95,255 | (38,83) | 16.30 [11.60, 21.01] |
| Is it OK for a person with HIV to teach | community dwelling women | 38,948 | (83) | 51.60% |
| Is it OK to care for a relative with HIV in household | community dwelling women | 38,948 | (83) | 63.50% |
| Is it ok to buy vegetables from a vendor with HIV? | community dwelling women | 38,948 | (83) | 41.10% |
| Is HIV transmission possible during pregnancy? | Students, general population(NDHS), Pregnant Women, community dwelling women | 96,498 | (14,16,38,60,83) | 45.09 [40.10, 50.08] |
| Is HIV transmission possible during delivery? | Students, general population, community dwelling women | 95,855 | (14,38,83) | 56.86 [54.43, 59.29] |
| Is HIV transmission possible during breastfeeding? | Students, general population, community dwelling women | 95,855 | (14,38,83) | 66.72 [62.89, 70.56] |
| Have you heard about HIV before? | Students, general population, Pregnant Women | 56,950 | (16,38,60) | 94.09 [91.18, 97.00] |
| Is HIV contracted through Sexual intercourse? | Students, Adolescents, FSW, Pregnant Women, | 2,446 | (14,16–18,23,48,60) | 68.09 [41.80, 94.38] |
| contracted through Sharing sharp unsterilized objects? | Students, FSW, Pregnant Women | 767 | (16,48,60) | 46.87 [10.49, 83.25] |
| Is HIV contracted through unscreened blood? | Students, FSW, Pregnant Women | 965 | (16,23,48,60) | 51.17 [4.52, 97.82] |
| Is HIV contracted through shaking hands? | Students, Dental tech students | 438 | (16,23) | 2.89 [-2.20, 7.98] |
| Is HIV contracted through witchcraft and other spiritual factors ? | general population(NDHS),community dwelling women | 95,255 | (38,83) | 24.05 [11.21, 36.89] |
| Is abstinence the best way of preventing HIV? | students, religious leaders | 776 | (23,63,81) | 55.77 [14.32, 97.22] |
| Is HIV contracted through intravenous needles? | Dental tech students, FSW, Pregnant Women | 725 | (23,48,60) | 61.47 [7.39, 115.56] |
| Is HIV contracted through sharing of cups/plates? | Students | 798 | (14,23) | 32.50 [-3.52, 68.51] |
| DO you knows a place to get HIV testing? | general population | 56,307 | (38) | 68.9 |

### South Africa

Nine included studies in regard to HIV were conducted among South Africans representing a total population of 17,320 participants; two studies were conducted in KwaZulu-Natal province ^[24,37]^ and one was conducted in each of Gauteng Province ^[29]^, Northern Cape province ^[42]^, North West province ^[34]^, two studies were toward online internet users ^[27,82]^, one study was conducted in Eastern Cape, Western Cape, Free State and Gauteng Provinces ^[46]^ while another study was nationally representative ^[84]^ (Table S4). The oldest among the study included was conducted in 2010 while the newest was conducted after 2010. Population under study was distributed among circumcised males, men who have sex with men or indicated interest in men, general population and home-based carers (Table S3). Majority of studies were conducted among both genders (5/9), while four were toward males only (ref). Age of respondents was from 15 to more than 25 years. Thirty two questions were asked to the participants that are related to the knowledge and awareness of HIV as general, transmission routes, clinical symptoms and prevention attitude, among which 16 questions were analyzed and synthesized. The question ‘Using condom will reduce HIV transmission?’’ was answered by 3,979 participants; 64.46% [95% Cl; 31.00, 97.91] answered yes. The question ‘’Do you perceive risk of contracting HIV?’’ was answered by 11,472 participants; 42.45% [95% Cl; −36.04, 120.05] answered yes. Questions asked, their corresponding study’s characteristics data, the pooled prevalence and the confidence intervals are depicted in (Table 4). Heterogeneity was high in all questions (I^2^ more than 80%).

**Table (4):**
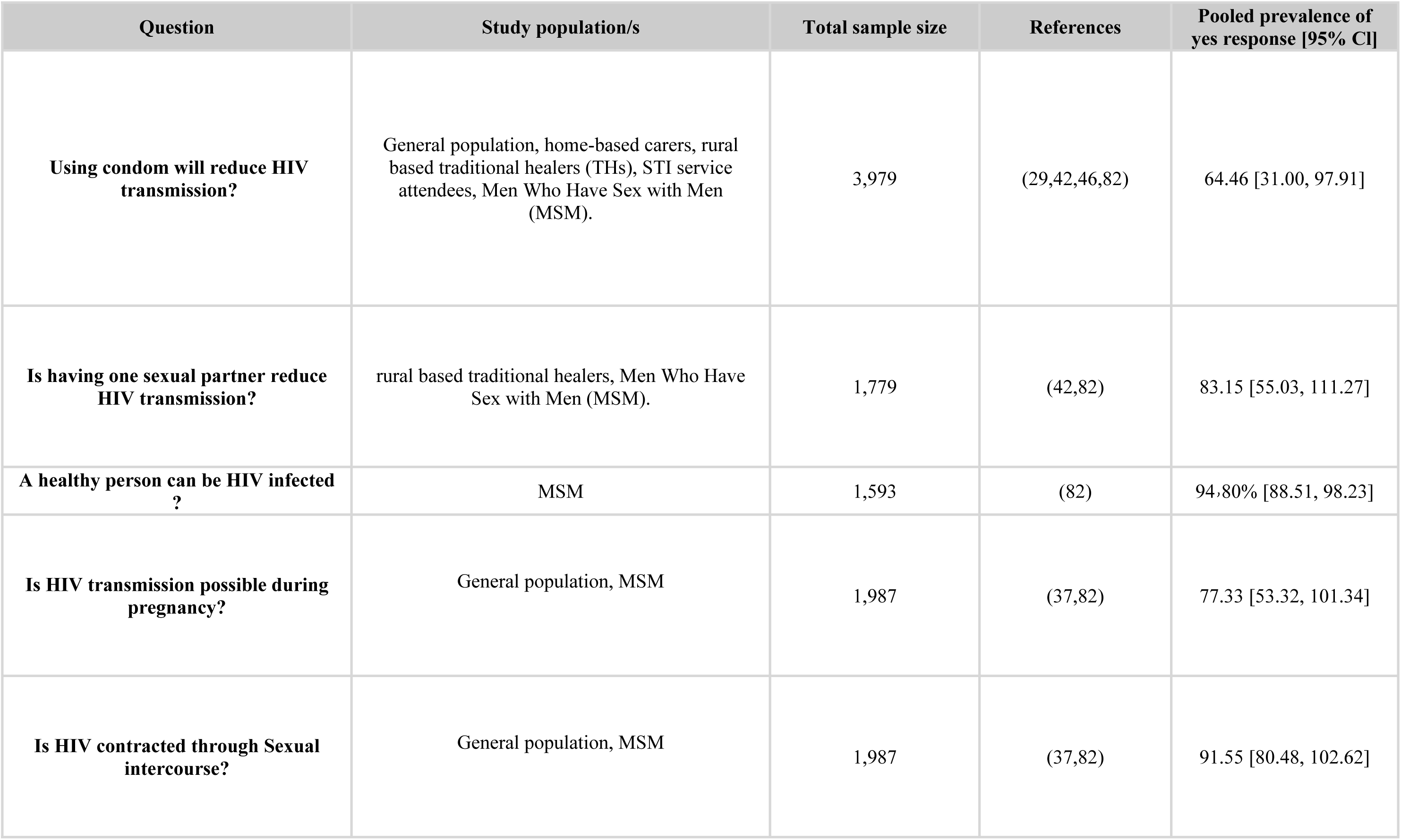
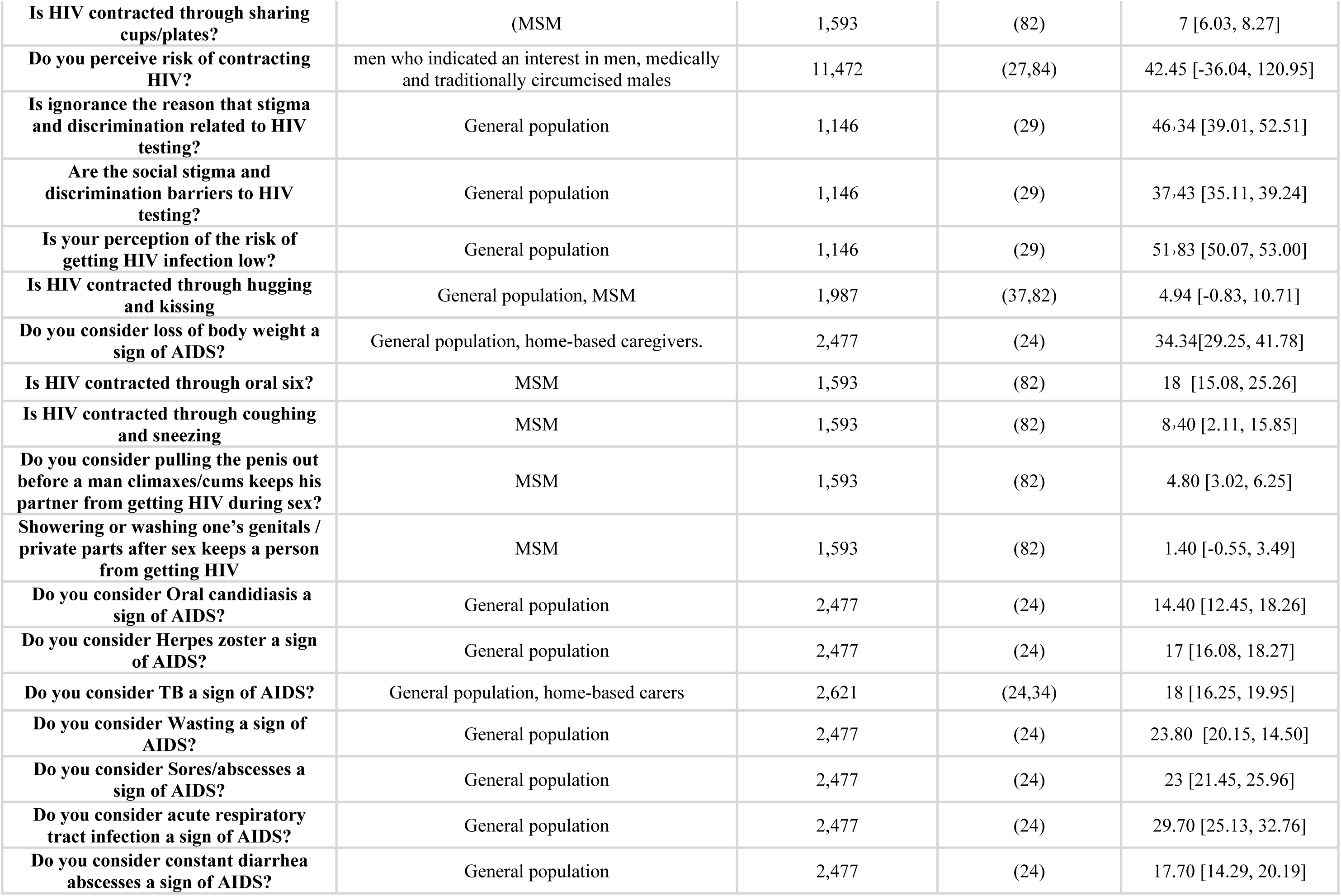
Awareness of HIV-related knowledge among South African population

### Adolescents

The study participants’ age were equal or less than 25 years in thirteen HIV-related included studies, representing a total population of 5,908 participants; five studies were conducted in Nigeria ^[14,16–18,63]^, three in Ghana ^[19,31,66]^, two in Mozambique ^[26,30]^, and one in each of Cameroon ^[59]^, Madagascar ^[71]^ and Uganda ^[72]^. Majority of studies were toward students and adolescents (11/13) while two studies were conducted among pregnant women. Majority of studies were conducted among both genders (11/13), while two were toward females only (pregnant women) (Table S5). Age of respondents was from 12 to 25 years. Twenty two questions were asked to the participants that are related to the knowledge and awareness of HIV as general, transmission routes, clinical symptoms and prevention attitude, among which 21 questions were analyzed and synthesized. The question ‘’Do you think HIV is contracted through Sexual intercourse?’’ was answered by 4,122 participants; 67.81% [95% Cl; 50.66, 84.96] answered yes. The question ‘’Do you think sharing cups/plates can transmit HIV?’’ was answered by 2,254 participants; 33.51% [95% Cl; 9.43, 57.59] answered yes. Questions asked, their corresponding studies’ characteristics, the pooled prevalence and the confidence intervals are depicted in (Table 5). Heterogeneity was high in all questions (I^2^ more than 80%).

**Table (5):**
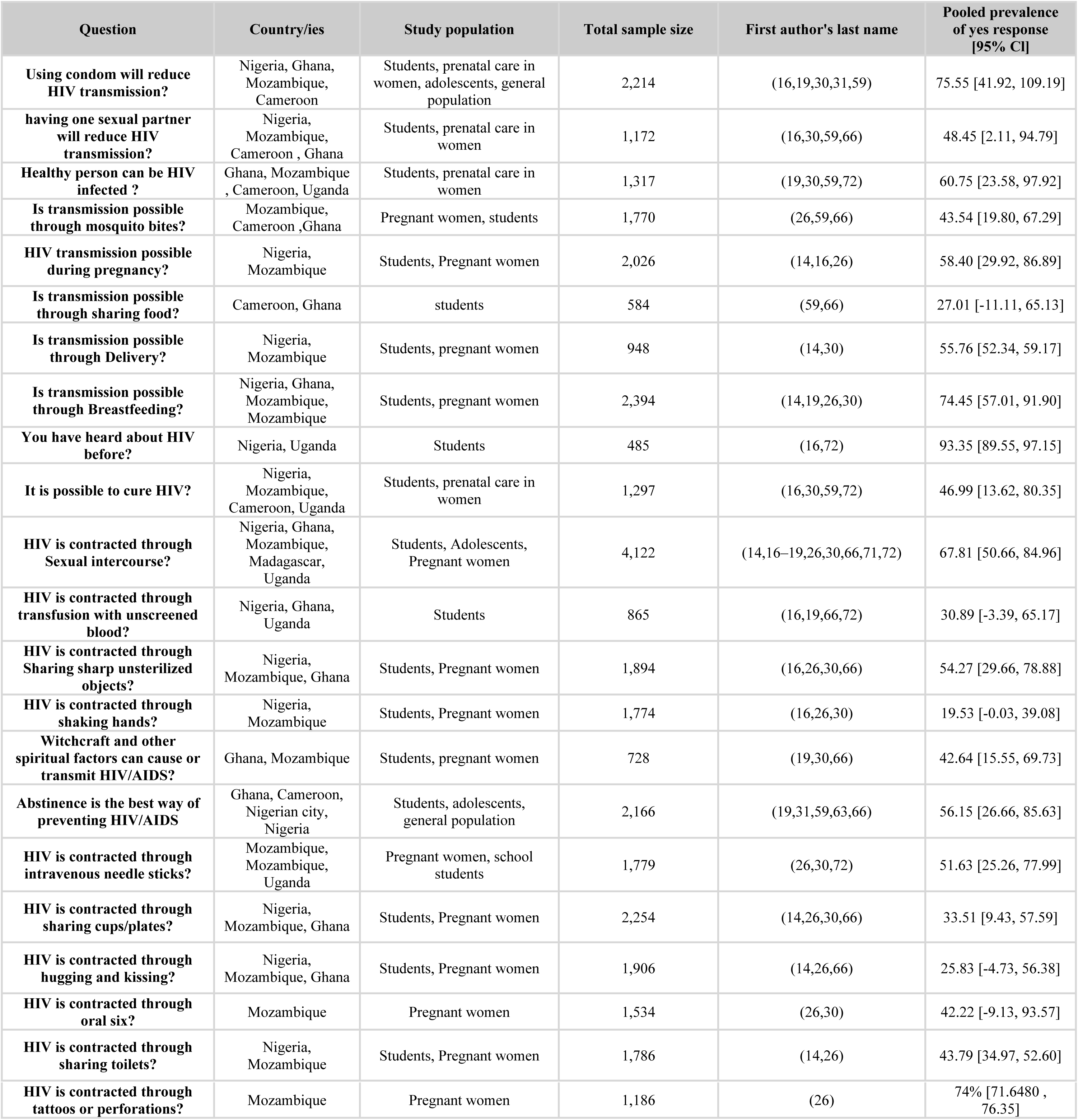
Awareness of HIV-related knowledge among adolescents in Africa

| Question | Country/ies | Study population | Total sample size | First author's last name | Pooled prevalence of yes response [95% CI] |
| --- | --- | --- | --- | --- | --- |
| Using condom will reduce HIV transmission? | Nigeria, Ghana, Mozambique, Cameroon | Students, prenatal care in women, adolescents, general population | 2,214 | (16,19,30,31,59) | 75.55 [41.92, 109.19] |
| having one sexual partner will reduce HIV transmission? | Nigeria, Mozambique, Cameroon , Ghana | Students, prenatal care in women | 1,172 | (16,30,59,66) | 48.45 [2.11, 94.79] |
| Healthy person can be HIV infected ? | Ghana, Mozambique , Cameroon, Uganda | Students, prenatal care in women | 1,317 | (19,30,59,72) | 60.75 [23.58, 97.92] |
| Is transmission possible through mosquito bites? | Mozambique, Cameroon ,Ghana | Pregnant women, students | 1,770 | (26,59,66) | 43.54 [19.80, 67.29] |
| HIV transmission possible during pregnancy? | Nigeria, Mozambique | Students, Pregnant women | 2,026 | (14,16,26) | 58.40 [29.92, 86.89] |
| Is transmission possible through sharing food? | Cameroon, Ghana | students | 584 | (59,66) | 27.01 [-11.11, 65.13] |
| Is transmission possible through Delivery? | Nigeria, Mozambique | Students, pregnant women | 948 | (14,30) | 55.76 [52.34, 59.17] |
| Is transmission possible through Breastfeeding? | Nigeria, Ghana, Mozambique, Mozambique | Students, pregnant women | 2,394 | (14,19,26,30) | 74.45 [57.01, 91.90] |
| You have heard about HIV before? | Nigeria, Uganda | Students | 485 | (16,72) | 93.35 [89.55, 97.15] |
| It is possible to cure HIV? | Nigeria, Mozambique, Cameroon, Uganda | Students, prenatal care in women | 1,297 | (16,30,59,72) | 46.99 [13.62, 80.35] |
| HIV is contracted through Sexual intercourse? | Nigeria, Ghana, Mozambique, Madagascar, Uganda | Students, Adolescents, Pregnant women | 4,122 | (14,16–19,26,30,66,71,72) | 67.81 [50.66, 84.96] |
| HIV is contracted through transfusion with unscreened blood? | Nigeria, Ghana, Uganda | Students | 865 | (16,19,66,72) | 30.89 [-3.39, 65.17] |
| HIV is contracted through Sharing sharp unsterilized objects? | Nigeria, Mozambique, Ghana | Students, Pregnant women | 1,894 | (16,26,30,66) | 54.27 [29.66, 78.88] |
| HIV is contracted through shaking hands? | Nigeria, Mozambique | Students, Pregnant women | 1,774 | (16,26,30) | 19.53 [-0.03, 39.08] |
| Witchcraft and other spiritual factors can cause or transmit HIV/AIDS? | Ghana, Mozambique | Students, pregnant women | 728 | (19,30,66) | 42.64 [15.55, 69.73] |
| Abstinence is the best way of preventing HIV/AIDS | Ghana, Cameroon, Nigerian city, Nigeria | Students, adolescents, general population | 2,166 | (19,31,59,63,66) | 56.15 [26.66, 85.63] |
| HIV is contracted through intravenous needle sticks? | Mozambique, Mozambique, Uganda | Pregnant women, school students | 1,779 | (26,30,72) | 51.63 [25.26, 77.99] |
| HIV is contracted through sharing cups/plates? | Nigeria, Mozambique, Ghana | Students, Pregnant women | 2,254 | (14,26,30,66) | 33.51 [9.43, 57.59] |
| HIV is contracted through hugging and kissing? | Nigeria, Mozambique, Ghana | Students, Pregnant women | 1,906 | (14,26,66) | 25.83 [-4.73, 56.38] |
| HIV is contracted through oral sex? | Mozambique | Pregnant women | 1,534 | (26,30) | 42.22 [-9.13, 93.57] |
| HIV is contracted through sharing toilets? | Nigeria, Mozambique | Students, Pregnant women | 1,786 | (14,26) | 43.79 [34.97, 52.60] |
| HIV is contracted through tattoos or perforations? | Mozambique | Pregnant women | 1,186 | (26) | 74% [71.6480 , 76.35] |

### Awareness related to demographic characteristics

Media (as general) was the main source of information of participants reported in several studies ^[17,47,74]^. However, other studies among students reported that school is the main source of information not media ^[22,75]^. Health professionals was the least mentioned source of information in the study of Saleh and colleagues ^[74]^.

Chaquisse and colleagues in their recently published study (2018) determined women’s age as not significantly associated with HIV and HBV knowledge. Moreover, they determined that to have heard about HIV/AIDS, Syphilis, Gonorrhoea, Hepatitis B or Hepatitis C, was associated with better knowledge about HIV transmission modes ^[26]^.

Two studies indicated a statistically significant difference in the HIV/AIDS knowledge scores and the marital/ relationship status ^[38,65]^. Nevertheless, another study indicated that no relation exists. This last study also reported that stigma toward HIV was significantly associated with knowledge scores of HIV as well as education level, female (sex) while place of residence (rural versus urban) is not^[55]^.

One study concluded that Comprehensive knowledge of HIV is significantly associated with more media items and fewer children at home ^[30]^.

Regarding religion, Christians compared to Muslims have been found to significantly have better knowledge of HIV/AIDS. Nevertheless, another study found that Muslim students scored higher on HIV/AIDS knowledge than Christian students ^[65,81]^.

Several studies indicated that the level of education and age have a significant association with the knowledge of HIV transmission ^[21,39,48]^. Additionally, one study ^[81]^ agreed that only education level is associated, while another agreed that only age is associated ^[77]^. Nevertheless, Faye and colleagues only concluded that marital status is associated to the knowledge of HIV transmission ^[39]^.

Seyoum and colleagues concluded that female participants who heard about HIV was significantly higher than that of the male participants. Moreover, there was a significant difference between males and females who suggested unsafe sexual intercourse as mode of transmission of HIV ^[77]^. However, Yaya and colleagues found that the majority of participants (females) (N=32,123, 82.5%) believe on contracting the virus via supernatural means as a mode of transmission ^[83]^.

### Hepatitis B Virus (HBV)

Fourteen included studies assessed the awareness of 9,446 Africans in regard to HBV, three studies were conducted in each of Nigeria ^[61,62,64]^, Cameroon ^[20,40,58]^ and Ghana ^[12,15,28]^, two in Ethiopia ^[32,52]^, one in each of Kenya, Mozambique and Madagascar ^[26,57,71]^. The oldest among the study included was conducted in 2010 while the newest was conducted in 2016 (Table 1). Population under study was found to be mainly students and adolescents and pregnant women (10/14), one study was targeting each of non medical staff of health facilities, general population, barbers and traders (Table 1). Majority of studies were conducted among both genders (8/14), five studies were toward females only (pregnant women) while one study included only males (barbers). Age of respondents ranged from 10 to 75 years. Fifteen questions were asked to the participants that are related to the knowledge and awareness of HBV as general, transmission routes, clinical symptoms, pathological consequences and prevention attitude, among which 13 questions were analyzed and synthesized. The question ‘’Do you know HBV?’’ was answered by 4,066 participants in Ghana, Mozambique, Ethiopia, Nigeria and Madagascar; 53.84% [95% Cl; 27.68, 79.99] answered yes. The question ‘’Does sexual contact is a possible route of HBV transmission?’’ was answered by 7,490 participants in Ghana, Mozambique, Ethiopia, Cameroon and Nigeria; 42.58% [95% Cl; 20.45, 64.71] answered yes. Questions asked, their corresponding articles’ data, the pooled prevalence and the confidence intervals are depicted in (Table 6). Heterogeneity was high in all questions (I^2^ more than 80%).

**Table (6):** Awareness of HBV-related knowledge among Africans

| Question | Country/ies | Study population/s | Total sample size | References | Pooled prevalence of yes response [95% CI] |
| --- | --- | --- | --- | --- | --- |
| **Do you know HBV?** | Ghana, Mozambique, Ethiopia, Kenya, Nigeria, Madagascar | Barbers, Pregnant women, students, general population | 4,570 | (12,15,26,28,52,57,62,71) | 53.84 [27.68, 79.99] |
| **HBV can damage liver?** | Ghana, Ethiopia, Cameroon, Nigeria | students, pregnant women, general population | 2,735 | (28,32,40,52,62) | 61.43 [31.34, 91.52] |
| **Does Blood transfusion is a possible route of HBV transmission?** | Ghana, Mozambique, Ethiopia, Cameroon, Nigeria | Barbers, Pregnant women, Students, traders, general population, Administration staff of a hospital | 7,490 | (15,26,32,40,52,58,61,62,64) | 56.16 [28.62, 83.69] |
| **Does reusing needles is a possible route of HBV transmission? (5)** | Ghana, Mozambique, Ethiopia, Cameroon, Nigeria | Barbers, Pregnant women, Students, general population, Administration staff of a hospital | 7,155 | (15,26,32,40,52,58,62,64) | 52.72 [26.96, 78.48] |
| Does sharing blades is a possible route of HBV transmission? | Ghana, Mozambique, Ethiopia, Cameroon, Nigeria | Barbers, Pregnant women, medicine and health science students, traders, Administration staff of a hospital | 5,437 | (15,26,32,40,61,64) | 39.75 [4.40, 75.10] |
| Does tattooing is a possible route of HBV transmission? | Ghana, Mozambique, Nigeria | Barbers, Pregnant women, traders, general population | 3,341 | (15,26,61,62) | 28.20 [5.03, 51.36] |
| Does sexual contact is a possible route of HBV transmission? | Ghana, Mozambique, Ethiopia, Cameroon, Nigeria | Barbers, Pregnant women, Students, traders, general population, Administration staff of a hospital | 7,490 | (15,26,32,40,52,58,61,62,64) | 42.58 [20.45, 64.71] |
| Does mosquito bites is a possible route of HBV transmission? | Mozambique, Nigeria | Pregnant women, general population | 2,806 | (26,62) | 28.49 [-3.85, 60.83] |
| Does mother to child is a possible route of HBV transmission? | Mozambique, Ethiopia, Nigeria | pregnant women, Students, general population | 3,745 | (26,28,32,52,62) | 57.55 [35.98, 79.12] |
| Hepatitis B can be transmitted through feco-oral route | Ethiopia, Cameroon | medical students | 519 | (32,58) | 48.65 [6.51, 90.79] |
| Does toothbrush sharing is a possible route of HBV transmission? | Nigeria | General population | 1,620 | (62) | 49 [ 46.6480 , 51.35] |
| HBV can be asymptomatic? | Ethiopia, Cameroon | pregnant women, medicine and health science students | 793 | (20,28,32,40) | 58.12 [23.38, 92.85] |
| HBV can be prevented by avoiding casual sex or multi sexual partnership? | Ethiopia | medicine and health science students | 408 | (32) | 87 [ 83.8641 , 90.13] |
| Do you know HBV vaccination? | Ethiopia, Cameroon | pregnant women, Students | 1,226 | (28,32,40,52,58) | 72.36 [50.69, 94.03] |

### Awareness related to demographic characteristics

Abdulai and colleagues in their study among pregnant women determined that level of education and occupation are significantly associated to hepatitis B awareness ^[12]^. Frambo and colleagues among the same population concluded that education is significantly associated to the level of awareness as well ^[40]^. Furthermore, Ngaira and colleagues assessed the awareness as well as vaccination status among the same population (pregnant women) and indicated a significant difference between vaccine uptake and education ^[57]^.

Noubiap and colleagues assessed HBV vaccine uptake but among medical students, and indicated that duration of study but not age and vaccination status are significantly correlated. Nevertheless Okonkwo and colleagues in their study among traders concluded that knowledge of the nature of HBV virus varied significantly according to age ^[58,61]^.

### Hepatitis C Virus (HCV)

Six included studies assessed the awareness of 2,306 Africans in regard to HCV, two studies were conducted in Egypt ^[74,79]^ and one in each of Ghana ^[15]^, Mozambique ^[26]^, Ethiopia ^[32]^ and Madagascar ^[71]^. The oldest among the study included was conducted after 2010 while the newest was conducted in 2015 (Table 1). Population under study was distributed among students and adolescents, general population, HCV positive patients, pregnant women and barbers (Table 1). Four studies were conducted among both genders, one toward females only and one toward males only (Table 1). Age of respondents range from 18 to 80 years. Seventeen questions were asked to the participants that are related to the knowledge and awareness of HCV as general, transmission routes, clinical symptoms, pathological consequences and prevention attitude, among which 10 questions were analyzed and synthesized. The question ‘’Does sexual contact is a possible route of HCV transmission?’’ was answered by 1,997 Africans in Ghana, Mozambique, Ethiopia and Egypt; 30.61% [95% Cl; 2.04, 59.17] answered yes. The question ‘’Hepatitis C infection can be prevented by vaccination?’’ was answered by 611 participants in Ethiopia and Egypt; 42.05 [95% Cl; 8.73, 75.37] answered yes. Questions asked, their corresponding articles’ data, the pooled prevalence and the confidence intervals are depicted in (Table 7). Heterogeneity was high in all questions (I^2^ more than 80%).

**Table (7):** Awareness of HCV-related knowledge among Africans

| Question | Country/ies | Study population/s | Total sample size | References | Pooled prevalence of yes response [95% CI] |
| --- | --- | --- | --- | --- | --- |
| Have you ever heard about viral hepatitis C? | Ghana, Mozambique, Egypt | Barbers, Pregnant women, university students, HCV patients | 1,831 | (15,26,71) | 20.21 [5.42, 35.00] |
| Do you know what cause HCV infection? | Egypt | General population | 67 | (74) | 87 [ 68.2002 , 87.79] |
| Is HCV can be cured? | Egypt | General population | 67 | (74) | 39 [ 27.4362 , 50.56] |
| Hepatitis C can cause liver cancer? | Ethiopia, Egypt | medicine and health science students, HCV patients | 611 | (32,79) | 89.50 [87.15, 91.85] |
| Does Blood transfusion is a possible route of HCV transmission? | Ghana | Barbers | 200 | (15,79) | 86.40 [81.70, 91.10] |
| Does reusing needles is a possible route of HCV transmission? | Ghana, Mozambique, Ethiopia, Egypt | Barbers, Pregnant women, medicine and health science students, HCV patients | 1,997 | (15,26,32,79) | 68.82 [16.96, 120.69] |
| Does sharing blades is a possible route of HCV transmission? | Ghana, Mozambique, Egypt | Barbers, Pregnant women, HCV patients | 1,589 | (15,26,79) | 54.72 [-9.47, 118.91] |
| Does tatooing is a possible route of HCV transmission? | Ghana, Mozambique | Barbers, Pregnant women | 1,386 | (15,26) | 21.60 [19.44, 23.76] |
| Does sexual contact is a possible route of HCV transmission? | Ghana, Mozambique, Ethiopia, Egypt | Barbers, Pregnant women, medicine and health science students, HCV patients | 1,997 | (15,26,32,79) | 30.61 [2.04, 59.17] |
| Can mother transmit HCV to infants? | Mozambique, Ethiopia | Pregnant women, medicine and health science students | 1,594 | (26,32) | 52.89 [-9.64, 115.41] |
| Does mosquito bite is a possible route of HCV transmission? | Mozambique | Pregnant women | 1,186 | (26) | 10.3 [ 8.7320 , 11.86] |
| Hepatitis C can be transmitted through feco-oral route? | Ethiopia | medicine and health science students | 408 | (32) | 70.3 [ 65.9881 , 74.61] |
| Does sharing toothbrush is a possible route of HCV transmission? | Mozambique, Egypt | Pregnant women, HCV patients | 1,389 | (32,79) | 52.46 [-7.32, 112.24] |
| Do you know any of HCV disease symptoms? | Egypt | Residents | 67 | (74) | 55 [ 43.2402 , 66.75] |
| infection can be asymptomatic? | Ethiopia | medicine and health science students | 408 | (32) | 55 [ 50.2961 , 59.70] |
| Hepatitis C infection can be prevented by vaccination? | Ethiopia, Egypt | medicine and health science students, HCV patients | 611 | (32,79) | 42.05 [8.73, 75.37] |
| Hepatitis C can be prevented by avoiding multi sexual partnership? | Ethiopia | medicine and health science students | 408 | (32) | 87 [ 83.8641 , 90.13] |

### Awareness related to demographic characteristics

Adoba and colleagues conducted their study among barbers - individuals of a sharp shaving objects’ based occupation, nevertheless, the radio was the major source of information on HCV infection (25.0 %)^[15]^.

Demsiss and colleagues in 2018 conducted a study among medicine and health science students in Ethiopia and determined that student’s residence and department were significantly associated with good level of knowledge toward transmission and prevention of hepatitis B and C infection ^[32]^.

### Human Papillomavirus (HPV)

Nine included studies assessed the awareness of 5,157 Africans in regard to HPV, four studies were conducted in Nigeria ^[35,36,41]^ and one in each of Madagascar, Morocco, Mali, South Africa and Senegal ^[43,49,51,54,70,71]^. The oldest among the study included was conducted in 2010 while the newest was conducted in 2016 (Table 1). Population under study was found to be mainly adolescents and students (6/9), while two studies was targeting general population and one was targeting HIV positive and negative females (Table 1). Majority of studies were conducted among both genders (6/9), while three studies were toward females only. Age of respondents range from 15 to older than 67 years. Fifteen questions were asked to the participants that are related to the knowledge and awareness of HPV as general, transmission routes, clinical symptoms, pathological consequences and prevention attitude, among which 13 questions were analyzed and synthesized. The question ‘’Do you know HPV?’’ was answered by 5,076 participants in Nigeria, Senegal, Morocco and Madagascar; 25.18% [95% Cl; 13.31, 37.06] answered yes. The question ‘’Are you aware of a vaccine for the prevention of HPV?’’ was answered by 2,548 participants in Nigeria and Morocco; 26.15% [95% Cl; 13.36, 38.93] answered yes. Furthermore; the question ‘’Do you know that HPV is a sexually transmitted infection’’ was answered by 1,409 participants in Nigeria, South Africa and Mali; 38.16% [95% Cl; 15.10, 61.22] answered yes. Questions asked, their corresponding articles’ data, the pooled prevalence and the confidence intervals are depicted in (Table 8). Heterogeneity was high in all questions (I^2^ more than 80%).

**Table (8):** Awareness of HPV-related knowledge among Africans

| Question | Country/ies | Study population/s | Total sample size | References | Pooled prevalence of yes response [95% CI] |
| --- | --- | --- | --- | --- | --- |
| Do you know HPV? | Nigeria; Senegal , Morocco , Madagascar | university staff and general population, mothers of female adolescents, students, Adolescents and young adults, parents | 5,076 | (35,36,41,49,51,54,71) | 25.18 [13.31, 37.06] |
| Do you know that HPV can cause cervical cancer | Nigeria, Mali | University students ,university staff and general population ,mothers of female adolescents ,adults and adolescents | 1,379 | (35,36,41,70) | 43.86 [19.39, 68.33] |
| HPV usually does not need treatment | South Africa | HIV positive and negative females | 30 | (43) | 97 [ 90.9241 , 103.075] |
| Cervical cancer is caused by persistent HPV infection | Nigeria | students | 368 | (49) | 19.6 [ 15.6801 , 23.51] |
| A person could have HPV without knowing it? | South Africa | HIV positive and negative females, university students | 310 | (41,43) | 79.92 [61.34, 98.51] |
| Vaccine may expose adolescents to risky sexual behaviors | Nigeria | mothers of female adolescents, university students | 570 | (36,41) | 34.65 [-20.82, 90.11] |
| having many sexual partners increases the risk of HPV? | South Africa, Nigeria | HIV positive and negative females, students | 678 | (41,43,49) | 65.64 [40.15, 91.13] |
| Do you know that HPV is a sexually transmitted infection | South Africa, Nigeria | University students ,university staff and general population , mothers of female adolescents ,HIV positive and negative females, adults and adolescents | 1,409 | (35,36,41,43,70) | 38.16 [15.10, 61.22] |
| Are you aware of a vaccine for the prevention of HPV? | Morocco, Nigeria | University students ,HIV positive and negative females, mothers of female adolescents, students, parents | 2,548 | (35,36,41,49,54) | 26.15 [13.36, 38.93] |
| Does not know where or how to access the vaccines | Nigeria | mothers of female adolescents, university students | 570 | (36,41) | 53.00 [48.98, 57.02] |
| The vaccine is too costly | Nigeria, South Africa | mothers of female adolescents ,HIV positive and negative females ,university students | 600 | (36,41,43) | 38.57 [13.65, 63.49] |
| Are you willing to receive the HPV vaccine? | Senegal, Mali, Nigeria | students , Adolescents and young adults | 2,985 | (41,49,51,70) | 58.54 [9.67, 107.41] |

### Awareness related to demographic characteristics

Funmilayo and colleagues in their study of medical students awareness and vaccination acceptance determined that the obstacles - as concluded by authors to receiving HPV vaccination among the female respondents included inadequate information (60.9%), high cost of vaccine (56.2%), poor access to vaccine (55.6%), worry about efficacy (38.5%), worry about safety (36.1%) and religious barriers (17.7%). A statistically significant association was found between level of awareness and vaccine acceptance and the level or class of students ^[41]^. Supporting this finding; Makwe and colleagues indicated the same association between the two variables as well ^[49]^.

Massey and colleagues in Senegal reported that respondents who indicated living most of their lives in a rural area demonstrated a greater percentage of ever having heard of HPV, and that fathers’ education level is significantly associated with the willingness of HPV vaccination. Mouallif and colleagues in Morocco concluded that mothers who agreed with the statement ‘Whatever happens to my health is God’s will’, believed that the vaccine was expensive and believed that they had insufficient information about the vaccine were significantly less likely to accept the vaccine ^[51,54]^.

### Sexually Transmitted Infections (STIs)

Seven included studies assessed the awareness of 2,986 Africans in regard to STIs as general, three studies were conducted in Nigeria ^[17,18,44]^ and one in each of Madagascar, Morocco, Mali, Uganda ^[47,56,70,71]^. The oldest among the study included was conducted after 2010 while the newest was conducted in 2014. Population under study was found to be mainly adolescents and students (5/7), while one study was targeting seafarers and another targeting women in reproductive age (Table 1). Majority of studies were conducted among both genders (5/7), one was toward males only while another was targeting females only (Table 1). Age of respondents range from 14 to older than 45 years. Thirty five questions were asked to the participants that are related to the knowledge and awareness of STIs general knowledge, transmission routes, clinical symptoms, pathological consequences and prevention attitude, among which 14 questions were analyzed and synthesized. The question ‘’Is Genital ulcer a symptom of having STIs?’’ was answered by 2,322 participants in Morocco and Uganda; 23.55% [95% Cl; 3.83, 43.27] answered yes. The question ‘’Do you know gonorrhea?’’ was answered by 1,123 participants in Nigeria and Madagascar; 22.84% [95% Cl; 5.13, 40.56] answered yes. questions asked, their corresponding articles’ data, the pooled prevalence and the confidence intervals are depicted in (Table 9). Heterogeneity was high in all questions (I^2^ more than 80%).

**Table (9):** Awareness of STIs-related knowledge among Africans

| Question | Country/ies | Study population/s | Total sample size | References | Pooled prevalence of yes response [95% CI] |
| --- | --- | --- | --- | --- | --- |
| Have you heard about STDs? | Nigeria | Adolescents, Students | 881 | (17,18) | 94.52 [89.62, 99.42] |
| **Do you know gonorrhea?** | Nigeria, Madagascar | Adolescents, Students, university students | 1,123 | (17,18,71) | 22.84 [5.13 , 40.56] |
| **Do you know syphilis?** | Nigeria, Madagascar | Students, university students | 782 | (18,71) | 5.67 [4.33, 7.01] |
| **Do you know Herpes simplex?** | Nigeria, Madagascar | Students, university students | 782 | (18,71) | 4.70 [1.17 , 8.23] |
| **Do you know chlamydia?** | Madagascar | university students | 242 | (71) | 1.70 [0.1320 , 3.268] |
| **Can Gonorrhea cause infertility?** | Nigeria | students | 30 | (44) | 60 [42.5563 , 77.4437] |
| **Is unprotected sex a mode of transmission of STIs?** | Nigeria | students | 540 | (18) | 87 [84.2561 , 89.7439] |
| **Is Needles and syringes a mode of transmission?** | Nigeria | students | 540 | (18) | 82.6 [79.4641, 85.7359] |
| **Is Blood and blood products a mode of transmission?** | Nigeria | students | 540 | (18) | 73.10 [69.3761, 76.823] |
| **Is Mother to child a mode of transmission of STIs?** | Nigeria | students | 540 | (18) | 70.9 [67.1761, 74.623] |
| **Is Coughing/sneezing a mode of transmission of STIs?** | Nigeria | students | 540 | (18) | 22 [18.6681, 25.3319] |
| **Is Sharing plates a mode of transmission of STIs?** | Nigeria | students | 540 | (18) | 12.2 [9.4561, 14.243] |
| **Can Gonorrhea transmits to neonates?** | Nigeria | students | 30 | (44) | 21.8 [6.5123 , 37.08] |
| **Is weight loss a symptom of having STIs?** | Nigeria, Uganda | Students, women of reproductive age | 875 | (18,56) | 41.44 [-28.24 , 111.11] |
| **Is Painful micturition a symptom of having STIs?** | Nigeria | students | 540 | (18) | 68.9 [65.1761, 72.62] |
| **Is Genital ulcer a symptom of having STIs?** | Nigeria, Uganda, Morocco | Students, women of reproductive age, seafarers | 2,322 | (18,47,56) | 23.55 [3.83 , 43.27] |
| **Is Genital swelling a symptom of having STIs?** | Nigeria | students | 540 | (18) | 38.4 [ 44.1841, 52.4159] |
| **Is Genital discharge a symptom of having STIs?** | Nigeria, Uganda, Morocco | Students, women of reproductive age, seafarers | 2,322 | (18,47,56) | 34.98 [-0.99, 70.95] |
| **Is Micturition burns a symptom of STIs in females?** | Morocco | seafarers | 1,447 | (47) | 6.2 [5.0240, 7.37] |
| **Is Micturition burns a symptom of STIs in males?** | Morocco | seafarers | 1,447 | (47) | 30.2 [ 27.7480, 32.45] |
| **Is Tumefaction of the groin a symptom of STIs in females?** | Morocco | seafarers | 1,447 | (47) | 1.2 [ 0.8080, 1.56] |
| **Is Tumefaction of the groin a symptom of STIs in males?** | Morocco | seafarers | 1,447 | (47) | 35 [ 32.6480, 37.35] |
| **Is Genital itching a symptom of STIs in females?** | Morocco, Uganda | seafarers, women of reproductive age | 1,782 | (47,56) | 34.09 [-16.28 , 84.75] |
| **Is Genital rash a symptom of STIs?** | Uganda | women of reproductive age | 335 | (56) | 14.5 [ 10.7761, 18.223] |
| **Is painful sex a symptom of STIs?** | Uganda | women of reproductive age | 335 | (56) | 0.3 [ -1.4640, 2.06] |
| **Can Gonorrhea be Asymptomatic ?** | Nigeria | students | 30 | (44) | 20.04 [ 5.5763, 34.58] |
| **Is abstinence a possible way to prevent STDs?** | Nigeria, Uganda | Adolescents, women of reproductive age | 676 | (17,56) | 23.20 [-19.23 , 85.63] |
| **Is condom use a possible way to prevent STDs?** | Nigeria, Uganda | Adolescents, women of reproductive age | 676 | (17,56) | 36.96 [13.44, 60.48] |
| **Being faithful by having one sexual partner is a way to prevent STIs?** | Uganda | women of reproductive age | 335 | (56) | 26.3 [ 21.5961 , 31.00] |
| **Do you use condoms?** | Nigeria | students | 30 | (44) | 72.5 [ 56.6243, 88.37] |
| **Do you reuse condoms?** | Nigeria | students | 30 | (44) | 10.1 [ -0.6798 , 20.87] |

### Awareness related to demographic characteristics

Akokuwebe and colleagues reported that Media (as general) was the main source of information 57% followed by friends 30%, and association between source of information about STDs is significantly related to age. Moreover, Laraqui and colleagues concluded that during the year prior to the study, 73.2% of participants (seafarers) were informed about the prevention of STI/HIV/AIDS through different ways, mainly the media (73% via TV and 45.6% via radio). Amu and colleagues provided more specific information in regard to source of knowledge as they determined that there are three major sources of information; the radio and television 343 (68.7%); teachers 340 (68.1%); and newspapers 224 (44.9%). Nevertheless, Nawagi and colleagues in their study in Uganda determined that only (23.9%) of the participants have information about STIs from the media ^[17,47,56]^.

Joda and his colleagues in Nigeria conducted a study to assess the level of knowledge of STIs among students from different schools and concluded that there is no statistically significant differences in the responses obtained from various schools. Moreover, Reuter and colleagues conducted a study to assess the difference of STIs related knowledge between university students of Madagascar and USA, and concluded that there is no statistically significant differences as well ^[44,71]^.

In spite of the study populations’ differences, five studies reported a significant association between knowledge of STIs and the level of education ^[30,38,68,81,82]^. Considering age as a factor affecting the level of awareness; several studies reported it to be significantly true ^[38,50,65,81]^, while other studies reported it is not ^[26,59]^. Moreover, living in an urban area was found to be significantly associated with awareness level in several studies ^[38,68,83]^.

## Discussion

The current study was the first of its kind - to our knowledge, as not general assessment of knowledge is studied, but the specific awareness determinants, the presented outcomes are believed to be the best inputs for organizing effective preventive measures as well as planning and conducting awareness raising campaigns.

The current study highlights the specific levels of STIs-related knowledge, practices and prevention attitudes among different African populations. The pooled prevalence estimates showed that even though more than 90% of the population had heard about STIs (94.52%) in general and HIV (92.24%) in particular, (79.79%) had never heard about HCV. These results are consistent with earlier studies in Eastern Europe, Victoria, Lao People’s Democratic Republic and Iran ^[85–88]^. Moreover, 25.18% of the population knows HPV. However, a study conducted among adolescents and adult women in one of the developed countries (USA) reported that only18% had heard about the virus ^[89].^ Nevertheless, the confounders among participants are to be considered when comparing the studies.

In the contrary of the expectation of HIV associated signs and symptoms awareness in such epidemic countries, this review revealed that almost only 14.4%, 17% and 17.7% of South Africans know that oral candidiasis, herpes zosters and constant diarrhea could be associated with HIV infection, respectively. Furthermore, less than 20% of the same population consider TB to be associated with AIDS ^[90]^.

The current findings of knowledge related to vertical HIV transmission during pregnancy 57%, delivery 66% or breastfeeding 73% corroborate with the studies of Stockholm et al (Stockholm 14) and Maimaiti et al (Maimaiti 15) although they slightly concluded higher proportions ^[91,92]^. Furthermore, these findings are in line with the results reported in UAE and Greece. Nevertheless, in India Pratibha Gupta and colleagues reported prevalence as low as 8.85% and 23.85% regarding the transmission during delivery and breastfeeding, respectively ^[93–95]^.

The findings clearly demonstrate that HIV preventative measures of South Africans are higher than that of Nigerians. For instance; using condom (64.46% versus 52.67%) and having one sexual partner (83.15% versus 57.67%) are known to reduce HIV transmission by South African and Nigerian populations, respectively. Bangladeshi women were reported to have knowledge similar to South Africans. However, other studies conducted in Vietnam, Italy and USA reported higher proportions [96–99].

It has been reported that increased HIV knowledge resulted in a reduction of risky sexual behaviors among adolescents ^[100]^. Notably, current findings revealed that adolescents in Africa were - for some extend aware of the facts associated with epidemics, transmission and prevention of HIV infection. Approximately 60.75% think that a healthy person can be HIV infected, similar finding was reported among Russians as well. Nevertheless, higher proportions were found to be reported in Iran and USA ^[101–103]^. More than 50% were found to be of good knowledge level about HIV transmission through Sexual intercourse (67.81%, Sharing sharp unsterilized objects (54.27%) and using intravenous needles (53.32%). This knowledge is higher when compared to Southern Brazilian adolescent’s. However, adolescents from India, US, Lao People’s Democratic Republic and Iraq were reported to possess higher knowledge scores ^[86,97,104–106]^.

Despite the finding that most of adolescents in Africa are aware of the correct ways of HIV transmission, they still express extensive misconceptions; nearly the half believe that HIV could be contracted through mosquitoes (43.54%), through toilet seats(43.79%), sharing cups/plates (33.51%) and through hugging or kissing (25.83%). Studies carried out among nursing students in Greece and among men who have sex with men in Finland illustrate similar findings as well. However, higher proportions (76%) for kissing and (100%) for sharing dishes, hot springs, kissing and mosquito bites were reported in Taiwan and Japan, respectively ^[95,107–109]^.

HIV-related stigma and discrimination persist as major obstacles to an effective HIV response in all parts of the world. Almost 37.4% of South Africans consider stigma is a barrier to HIV testing. Generally speaking, Africans’ attitudes towards HIV/AIDS patients were in need for enforcement. For example, 62.95% would care for a relative with HIV in household, 57.11% would buy vegetables from an HIV infected vendor, and only 44.80% would allow a person with HIV to teach. Similar results were found to be reported in Sri Lanka. However, Janahi and colleagues in their findings reported that more than half of the adult participants (n=1,630) in Bahrain would avoid sitting near, hugging or even shaking HIV infected people hands ^[110–112]^.

The findings presented in this study regarding HBV illustrate that the knowledge of Africans is moderate, i.e. 61.43% know about the consequence of liver damage. Moreover, reusing needles 52.72%, sexual contact 42.58% and toothbrush sharing 49% were considered to be possible routs of HBV transmission. Furthermore, 72.36% correctly believe in the existence of vaccination. A prior study conducted among Asian Americans in USA reported similar or slightly higher knowledge ^[113]^.

Regarding HCV, about 68.82% of Africans have correct knowledge about its transmission through reusing needles. However, nearly the half (42.05%) incorrectly believe in the existence of vaccination. Taiwanese dental students also believe that there is an effective vaccine for HCV but in a proportion as low as 15% ^[107]^.

The pooled prevalence of the African knowledge regarding the association between HPV and cervical cancer was found to be mostly 43.86%, this is consistent with a study conducted in USA. Moreover, nearly 26.15% of South Africans were aware of a vaccine for HPV prevention, lower prevalence (10.8%, N=1,177) was reported in Berlin, Germany recently (2018). However, the fact that the later study was conducted among students and young adults is needed to considered when comparing the results. ^[114,115]^.

Implementation of educational awareness programs in schools will have its impacts in the near future. Moreover, knowledge raising campaigns at the continent level or nationally, in urban and rural regions targeting infected or non infected individuals using traditional sittings or integrating new online tools is needed to be applied, for raising awareness, increasing willingness for testing, decreasing STIs transmission and decreasing stigma and discriminations.

### Strengths and Limitations

The strengths of this review are that we systematically identified and included awareness estimates from 2010, Moreover, we have conducted meta-analysis to derive pooled prevalence estimates of all questions related. Furthermore, we carried out a quality assessment of the included studies based on criteria specifically developed to determine the quality of included studies.

Nevertheless, several limitations are to be considered when interpreting study results; grey literature evidence was not assessed. Moreover, African journals that are not indexed in the screened databases was not considered for inclusion as well, although all included studies are of good quality, several good studies might have be missed. Furthermore, another parameter that should be considered is that the limited number of participants in some questions can be observed for which the outcome might not be suitable to be generalized to the continent/country/population level. Lastly, the heterogeneity was high among the majority of questions analyzed.

## Conclusion

The current study indicates that awareness is needed to be enforced. The differences observed among populations are highlighting the possibility for containment and control by directing light toward specific populations or countries as well as addressing specific awareness knowledge to ensure that the general as well as the related specific preventive awareness knowledge is improved.

## Authors’ contribution

Designed the experiments: BMM and AMS. Performed the experiments: All authors. Analyzed the data: BMM. Wrote the paper: BMM, AMS and OWM. All authors read and approved the final article.

---

**Table (S1):** PRISMA checklist.

| Section/topic | # | Checklist item | Reported on page # |
| --- | --- | --- | --- |
| **TITLE** |  |  |  |
| Title | 1 | Identify the report as a systematic review, meta-analysis, or both. | 1 |
| **ABSTRACT** |  |  |  |
| Structured summary | 2 | Provide a structured summary including, as applicable: background; objectives; data sources; study eligibility criteria, participants, and interventions; study appraisal and synthesis methods; results; limitations; conclusions and implications of key findings; systematic review registration number. | 1 |
| **INTRODUCTION** |  |  |  |
| Rationale | 3 | Describe the rationale for the review in the context of what is already known. | 2 |
| Objectives | 4 | Provide an explicit statement of questions being addressed with reference to participants, interventions, comparisons, outcomes, and study design (PICOS). | 2 |
| **METHODS** |  |  |  |
| Protocol and registration | 5 | Indicate if a review protocol exists, if and where it can be accessed (e.g., Web address), and, if available, provide registration information including registration number. | 3 |
| Eligibility criteria | 6 | Specify study characteristics (e.g., PICOS, length of follow-up) and report characteristics (e.g., years considered, language, publication status) used as criteria for eligibility, giving rationale. | 3 |
| Information sources | 7 | Describe all information sources (e.g., databases with dates of coverage, contact with study authors to identify additional studies) in the search and date last searched. | 3 |
| Search | 8 | Present full electronic search strategy for at least one database, including any limits used, such that it could be repeated. | 3 |
| Study selection | 9 | State the process for selecting studies (i.e., screening, eligibility, included in systematic review, and, if applicable, included in the meta-analysis). | 4 |
| Data collection process | 10 | Describe method of data extraction from reports (e.g., piloted forms, independently, in duplicate) and any processes for obtaining and confirming data from investigators. | 4 |
| Data items | 11 | List and define all variables for which data were sought (e.g., PICOS, funding sources) and any assumptions and simplifications made. | 4 |
| Risk of bias in individual studies | 12 | Describe methods used for assessing risk of bias of individual studies (including specification of whether this was done at the study or outcome level), and how this information is to be used in any data synthesis. | 4 |
| Summary measures | 13 | State the principal summary measures (e.g., risk ratio, difference in means). | 4 |
| Synthesis of results | 14 | Describe the methods of handling data and combining results of studies, if done, including measures of consistency (e.g., I<sup>2</sup>) for each meta-analysis. | 5 |
| Risk of bias across studies | 15 | Specify any assessment of risk of bias that may affect the cumulative evidence (e.g., publication bias, selective reporting within studies). | 4 |
| Additional analyses | 16 | Describe methods of additional analyses (e.g., sensitivity or subgroup analyses, meta-regression), if done, indicating which were pre-specified. | 5 |
| **RESULTS** |  |  |  |
| Study selection | 17 | Give numbers of studies screened, assessed for eligibility, and included in the review, with reasons for exclusions at each stage, ideally with a flow diagram. | 5 |
| Study characteristics | 18 | For each study, present characteristics for which data were extracted (e.g., study size, PICOS, follow-up period) and provide the citations. | 6 |
| Risk of bias within studies | 19 | Present data on risk of bias of each study and, if available, any outcome level assessment (see item 12). | 9 |
| Results of individual studies | 20 | For all outcomes considered (benefits or harms), present, for each study: (a) simple summary data for each intervention group (b) effect estimates and confidence intervals, ideally with a forest plot. | 11 |
| Synthesis of results | 21 | Present results of each meta-analysis done, including confidence intervals and measures of consistency. | 11 |
| Risk of bias across studies | 22 | Present results of any assessment of risk of bias across studies (see Item 15). | 8 |
| Additional analysis | 23 | Give results of additional analyses, if done (e.g., sensitivity or subgroup analyses, meta-regression [see Item 16]). | 11 |
| **DISCUSSION** |  |  |  |
| Summary of evidence | 24 | Summarize the main findings including the strength of evidence for each main outcome; consider their relevance to key groups (e.g., healthcare providers, users, and policy makers). | 28 |
| Limitations | 25 | Discuss limitations at study and outcome level (e.g., risk of bias), and at review-level (e.g., incomplete retrieval of identified research, reporting bias). | 30 |
| Conclusions | 26 | Provide a general interpretation of the results in the context of other evidence, and implications for future research. | 30 |
| **FUNDING** |  |  |  |
| Funding | 27 | Describe sources of funding for the systematic review and other support (e.g., supply of data); role of funders for the systematic review. | 30 |

**Table (S2):** Assessment of quality of included studies.

| Study | Was the study obj clear? | Study sample clearly defined | Study pop specified and defined? | Participation rate above 70%? | Sitting and methods rigorous? | Analysis rigorous? | Total score |
| --- | --- | --- | --- | --- | --- | --- | --- |
| Abdulai *et al* (12) | 1 | 1 | 1 | 0 | 1 | 1 | 5 |
| Abimanyi-Ochom *et al* (13) | 1 | 1 | 0 | 0 | 1 | 1 | 4 |
| Aderemi *et al* (14) | 1 | 1 | 1 | 0 | 1 | 1 | 5 |
| Adoba *et al* (15) | 1 | 1 | 1 | 0 | 0 | 1 | 4 |
| Ajide and Balogun (16) | 1 | 1 | 1 | 0 | 0 | 1 | 4 |
| Akokuwebe *et al* (17) | 1 | 1 | 1 | 0 | 0 | 1 | 4 |
| Amu and Adegun (18) | 1 | 1 | 1 | 1 | 0 | 1 | 5 |
| Appiah-Agyekum *et al* (19) | 1 | 0 | 1 | 0 | 1 | 1 | 4 |
| Aroke *et al* (20) | 1 | 1 | 1 | 0 | 0 | 1 | 4 |
| Asefa and Beyene (21) | 1 | 1 | 0 | 1 | 1 | 1 | 5 |
| Audet *et al*(22) | 1 | 1 | 1 | 0 | 0 | 1 | 4 |
| Azodo *et al* (23) | 1 | 1 | 1 | 1 | 0 | 1 | 5 |
| Becker *et al* (24) | 1 | 1 | 0 | 0 | 1 | 0 | 3 |
| Carlos *et al* (25) | 1 | 1 | 1 | 0 | 0 | 1 | 4 |
| Chan and Tsai (8) | 1 | 1 | 0 | 1 | 0 | 1 | 4 |
| Chaquisse *et al* (26) | 1 | 1 | 0 | 1 | 1 | 1 | 5 |
| Chard *et al* (27) | 1 | 1 | 1 | 1 | 0 | 1 | 5 |
| Cheng *et al* (28) | 1 | 1 | 1 | 0 | 0 | 1 | 4 |
| Chimoyi *et al* (29) | 1 | 0 | 1 | 0 | 0 | 1 | 3 |
| Ciampa *et al* (30) | 1 | 1 | 0 | 1 | 1 | 1 | 5 |
| Darteh *et al* (31) | 1 | 1 | 0 | 1 | 0 | 1 | 4 |
| Demsiss *et al* (32) | - | - | - | - | - | - | - |
| Elbadawi *et al* (33) | 1 | 1 | 1 | 1 | 0 | 1 | 5 |
| Engelbrecht *et al* (34) | 1 | 1 | 1 | 0 | 0 | 1 | 4 |
| Eni *et al* (35) | 1 | 1 | 0 | 0 | 1 | 1 | 4 |
| Ezenwa *et al* (36) | 1 | 1 | 1 | 1 | 0 | 1 | 5 |
| Faleye *et al* (37) | 1 | 1 | 1 | 1 | 0 | 1 | 5 |
| Faust *et al* (38) | 1 | 1 | 0 | 0 | 0 | 1 | 3 |
| Faye *et al* (39) | 1 | 1 | 1 | 1 | 1 | 1 | 6 |
| Frambo *et al* (40) | 1 | 1 | 0 | 1 | 0 | 1 | 4 |
| Funmilayo *et al* (41) | 1 | 1 | 0 | 0 | 0 | 1 | 3 |
| George *et al* (42) | 1 | 1 | 0 | 1 | 0 | 0 | 3 |
| Griffith *et al* (43) | 1 | 1 | 0 | 1 | 0 | 0 | 3 |
| Joda *et al*(44) | 1 | 1 | 0 | 1 | 0 | 0 | 3 |
| Kiderlen *et al*(45) | 1 | 1 | 1 | 0 | 0 | 1 | 4 |
| Kufa *et al* (46) | 1 | 1 | 0 | 1 | 0 | 1 | 4 |
| Laraqui *et al* (47) | 1 | 1 | 1 | 1 | 0 | 1 | 5 |
| Lawan *et al* (48) | 1 | 1 | 0 | 0 | 1 | 1 | 4 |
| Makwe *et al* (49) | 1 | 1 | 1 | 1 | 0 | 1 | 5 |
| Mason *et al* (50) | 1 | 1 | 1 | 0 | 0 | 1 | 4 |
| Massey *et al*(51) | 1 | 1 | 1 | 1 | 0 | 1 | 5 |
| Mesfin *et al* (52) | 1 | 1 | 1 | 0 | 0 | 1 | 4 |
| Mkumbo *et al* (53) | 1 | 1 | 1 | 1 | 0 | 1 | 5 |
| Mouallif *et al* (54) | 1 | 1 | 1 | 0 | 0 | 1 | 4 |
| Nabukenya *et al* (55) | 1 | 1 | 0 | 1 | 0 | 1 | 4 |
| Nawagi *etal* (56) | 1 | 1 | 1 | 0 | 0 | 1 | 4 |
| Ngaira *et al* (57) | 1 | 1 | 1 | 0 | 1 | 0 | 4 |
| Noubiap *et al* (58) | 1 | 1 | 1 | 0 | 1 | 0 | 4 |
| Nubed *et al* (59) | 1 | 1 | 0 | 1 | 0 | 1 | 4 |
| Ojieabu *et al* (60) | 1 | 1 | 1 | 1 | 0 | 1 | 5 |
| Okonkwo *et al* (61) | 1 | 1 | 0 | 1 | 0 | 0 | 3 |
| Okonkwo *et al* (62) | 1 | 1 | 1 | 1 | 1 | 1 | 6 |
| Oladebo *and* Fayemi (63) | 1 | 1 | 1 | 0 | 1 | 0 | 4 |
| Omotowo *et al*(64) | 1 | 1 | 1 | 1 | 1 | 1 | 6 |
| Oppong and Oti-Boadi (65) | 1 | 1 | 1 | 0 | 0 | 0 | 3 |
| Owusu (66) | 1 | 0 | 1 | 0 | 0 | 1 | 3 |
| Oyekale (67) | 1 | 1 | 0 | 0 | 1 | 1 | 4 |
| Paintsil *et al* (68) | 1 | 1 | 1 | 0 | 0 | 1 | 4 |
| Pathmanathan *et al*(69) | 1 | 1 | 0 | 0 | 0 | 1 | 3 |
| Poole *et al* (70) | 1 | 1 | 1 | 0 | 1 | 1 | 5 |
| Reuter *et al* (71) | 1 | 0 | 1 | 0 | 1 | 1 | 4 |
| Rukundo *et al* (72) | 1 | 1 | 0 | 0 | 1 | 1 | 4 |
| Sahile *et al* (73) | 1 | 1 | 1 | 1 | 0 | 1 | 5 |
| Saleh *et al* (74) | 1 | 1 | 0 | 0 | 1 | 1 | 4 |
| Sandqvist *et al* (75) | 1 | 1 | 1 | 0 | 1 | 0 | 4 |
| Schwitters *et al* (76) | 1 | 1 | 1 | 0 | 1 | 1 | 5 |
| Seyoum and Legesse (77) | 1 | 1 | 1 | 0 | 1 | 1 | 5 |
| Shiferaw *et al* (78) | 1 | 1 | 1 | 0 | 0 | 1 | 4 |
| Sultan *et al* (79) | 1 | 1 | 1 | 0 | 0 | 1 | 4 |
| Tarekegne *et al* (80) | 1 | 1 | 1 | 1 | 1 | 1 | 6 |
| Umar and Oche (81) | 1 | 1 | 1 | 1 | 0 | 1 | 5 |
| Wagenaar *et al* (82) | 1 | 0 | 1 | 0 | 0 | 1 | 3 |
| Yaya *et al* (83) | 1 | 1 | 0 | 1 | 1 | 1 | 5 |
| Zungu *et al* (84) | 1 | 1 | 1 | 0 | 1 | 1 | 5 |

**Table (S3):** Characteristics of HIV-related included studies conducted among Nigerians.

| Study | Year of publication | Year/s of conduction | City/Region | study population/s | sample size | Gender | Participants' Age |
| --- | --- | --- | --- | --- | --- | --- | --- |
| Aderemi *et al* (14) | 2013 | After 2010 | Oyo State | Students | 600 | Both | 12-19 |
| Ajide and Balogun (16) | 2018 | After 2010 | Ibadan/Oyo State | Students | 240 | Both | 16±1 |
| Akokuwebe *et al* (17) | 2016 | After 2010 | Ikeji-Arakeji/Osun State | Adolescents | 341 | Both | 16+-2 |
| Amu and Adegun (18) | 2015 | After 2010 | Ekiti State | Students | 540 | Both | Oct-14 |
| Azodo *et al* (23) | 2014 | 2010 | Enugu State | Dental tech students | 198 | Both | 20- ≥27 |
| Faust *et al* (38) | 2017 | 2013 | Nigeria | general population | 56 307 | Both | 15–49 |
| Lawan *et al*(48) | 2012 | 2011 | KanoState | FSW | 124 | Females | 26±2 |
| Ojieabu *et al* (60) | 2011 | 2011 | Sagamu/Ogun State | Pregnant Women | 403 | Females | 20- ≥40 |
| Oladebo and Fayemi (63) | 2011 | 2010 | Ibadan/Oyo State | Secondary students | 420 | Both | 10-19 |
| Umar and Oche (81) | 2012 | 2010 | Sokoto State | religious leaders | 158 | Male | ≥30 |
| Yaya *et al* (83) | 2018 | 2013 | Nigeria | community dwelling women | 38 948 | Females | 15-49 |

**Table (S4):** Characteristics of HIV-related included studies conducted among South Africans.

| Study | Year of publication | Year/s of conduction | City/Region | study population/s | sample size | Gender | Participants' Age |
| --- | --- | --- | --- | --- | --- | --- | --- |
| Becker *et al*(24) | 2015 | 2009-2010 | Durban/KwaZulu-Natal | General population | 2,477 | Both | N.A |
| Chard *et al* (27) | 2017 | After 2010 | South Africa | men who indicated an interest in men | 386 | Male | mean 33 |
| Chimoyi *et al* (29) | 2015 | 2013 | Johannesburg/Gauteng | General population | 1,146 | Both | N.A |
| Engelbrecht *et al* (34) | 2017 | 2012 | Moretele Sub-district/Bojanala District/North West province | home-based carers | 144 | Both | median 35 |
| Faleye *et al* (37) | 2014 | 2012-2013 | Durban/KwaZulu-Natal/South africa | Male medical circumcision clients | 394 | Male | mean 28±9 |
| George *et al* (42) | 2013 | 2010 | Northern Cape province | rural based traditional healers | 186 | Both | N.A |
| Kufa *et al* (46) | 2018 | After 2010 | Eastern Cape,Western Cape,Free State,Gauteng State | STI service attendees | 1,054 | Both | 23-32 |
| Wagenaar *et al* (82) | 2012 | 2010 | South Africa | Men Who Have Sex with Men | 1,593 | Male | ≥18 |
| Zungu *et al* (84) | 2016 | 2012 | South Africa | medically and traditionally circumcised males | 11,086 | Male | ≥15 |

**Table (S5):**
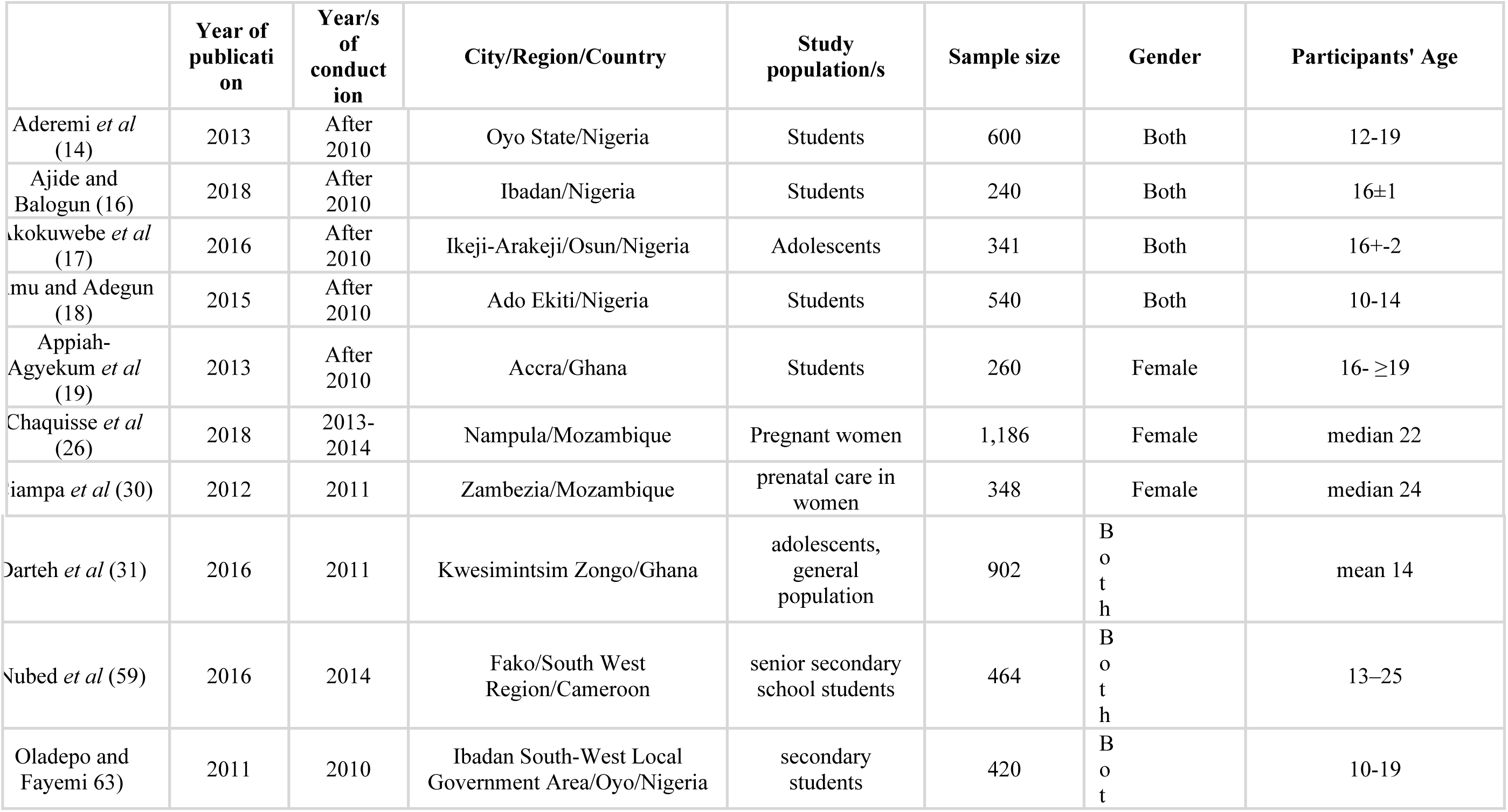
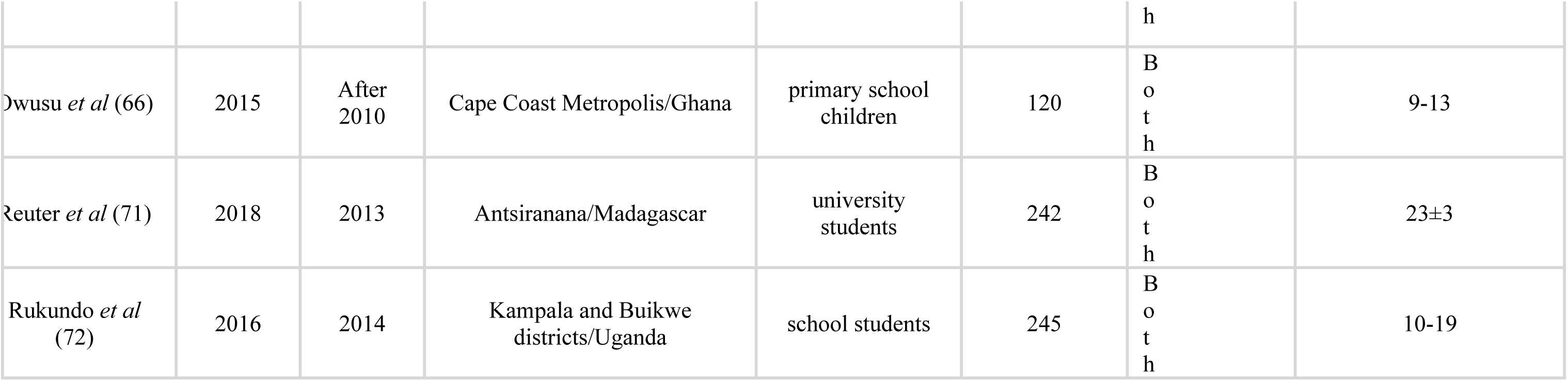
Characteristics of HIV-related included studies conducted among adolescent Africans.

